# Stoichiometric m6A regulation of YY1 orchestrates the metabolic switch during cardiac development

**DOI:** 10.64898/2026.09.25.754444

**Authors:** Shunyao Lei, Harshi Gangrade, Sheetal Bajpayi, Qingqing Gong, Courtney Hall, Edwin Jong-woo Yoo, Simone Sidoli, Nanami Senoo, Konstantinos Boulias, Steven M. Claypool, Winston Timp, Emmanouil Tampakakis

## Abstract

Chemical modifications of RNA provide an additional regulatory layer essential for development. N6-methyladenosine (m6A) is the most abundant internal mRNA mark, but how site specificity, stoichiometry, and cellular context together determine mRNA fate remains unclear. Here, we define these rules in cardiogenesis by combining single-molecule nanopore direct RNA sequencing (DRS) with a CRISPR-dCas13 epitranscriptomic editing platform in human induced pluripotent stem cells and a conditional Mettl14 cardiac knockout in mice, integrated with multi-omics, to map m6A sites and quantify stoichiometry. We uncover a bimodal m6A code in which low versus high modification ratios are associated with opposite mRNA and protein outcomes. We further show that Mettl14 loss causes developmental cardiomyopathy and embryonic lethality by disrupting the glycolysis-to-oxidative phosphorylation (OXPHOS) metabolic switch, and identify an m6A–YY1 (Yin Yang 1) axis that post-transcriptionally controls OXPHOS genes. Finally, targeted installation of m6A at defined sites using CRISPR–dCas13 reveals that m6A effects on YY1 are strongly locus- and context-dependent. Together, these findings establish m6A as a site- and stoichiometry-dependent regulatory code interpreted by cellular context to direct cardiac metabolic maturation. These principles are likely to extend to other developmental and physiological contexts in which precise control of metabolic state and gene dosage is required.

## Introduction

The precise temporal and spatial regulation of gene expression underlies multicellular development, dictating cell fate transitions, metabolic adaptation, and the formation of complex organs. While classical genetic and transcriptional mechanisms are well-understood, many developmental processes and the underlying causes of diseases are still not fully explained, highlighting the need for understanding additional layers of regulatory control^1–4^. Among these, epitranscriptomics – the dynamic and reversible chemical modification of RNA – has emerged as an important yet mechanistically underexplored regulatory layer that shapes gene expression landscapes and cellular programs^5–7^. However, the exact molecular rules governing how these modifications precisely dictate cellular outcomes, especially during rapid developmental transitions and the establishment of functional organ systems, remain a significant frontier in molecular and developmental biology.

N6-methyladenosine (m6A) is the most abundant internal mRNA modification, ubiquitously influencing all stages of mRNA biology, from splicing and nuclear export to stability and translation^8–11^. This dynamic mark is deposited by ‘writers’ such as methyltransferase-like 3 (Mettl3) and methyltransferase-like 14 (Mettl14) that form a complex with Wilms’ tumor-1 associated protein (WTAP), removed by ‘erasers’ such as AlkB homolog-5 (Alkbh5) and alpha ketoglutarate-dependent dioxygenase (Fto) and interpreted by diverse ‘readers’ that are mainly YTH domain-containing proteins (YTHDC1, YTHDF1-3) ^8,10–12^. While m6A is known to be context-dependent and frequently found in conserved DRACH motifs^8,10,12^, a profound knowledge gap persists in understanding the fundamental molecular logic of the m6A code. Specifically, it is unclear how the stoichiometric ratio of m6A and its site-specific placement across transcripts translates into distinct, and potentially divergent, downstream regulatory effects that ultimately control gene expression. Conventional methodologies, often relying on antibody-based enrichment and RNA fragmentation, critically lack the single-molecule, full-length transcript resolution necessary to decipher these nuanced site-specific and stoichiometric effects, hindering a comprehensive mechanistic interpretation of m6A’s impact on mRNA fate and its dynamic interplay with cellular processes^13–15^.

Despite the pervasive regulatory potential of m6A epitranscriptomics across cell biology^16–20^, its precise orchestrating mechanisms during dynamic cellular processes like early organogenesis – a period characterized by intense metabolic and morphological remodeling – remain largely unexplored. Previous studies have highlighted m6A’s importance in stem cell identity, differentiation potential, and pathological conditions such as impaired human stem cell-derived cardiomyocytes (hiPSC-CMs) differentiation and cardiac diseases^11,21–28^. However, a critical unmet need is a high-resolution, mechanistic understanding of how m6A fundamentally controls these rapid and profound changes during such crucial developmental windows, particularly in complex organs like the developing heart, where post-transcriptional pathways are upregulated^17,18,29,30^. Furthermore, the question of whether m6A’s effects are uniformly applied or exquisitely context-dependent (i.e., cell-type or developmental stage-specific) represents a major unresolved issue in the field.

Here, we investigate how m6A epitranscriptomics governs developmental gene expression and metabolism, using cardiac formation as a model for dynamic cell-fate and metabolic transitions. Through high-depth nanopore direct RNA sequencing (DRS), we generate a comprehensive, site-specific m6A landscape and quantitative stoichiometry map of mouse embryonic hearts. This single-molecule approach reveals a bimodal m6A regulatory code in which distinct modification ratios on individual transcripts are associated with opposite effects on mRNA and protein abundance, challenging the notion of a uniform m6A outcome. We demonstrate that conditional deletion of the m6A writer Mettl14 results in severe developmental cardiomyopathy and embryonic lethality by disrupting the metabolic switch from glycolysis to oxidative phosphorylation (OXPHOS). Integrating our multi-omics analyses, we further uncover a post-transcriptional m6A–YY1 regulatory axis that orchestrates this metabolic transition by controlling OXPHOS gene expression via the transcription factor Yin Yang 1 (YY1). Finally, we develop a CRISPR-based hiPSC platform for targeted epitranscriptomic editing, which reveals that m6A-mediated regulation of YY1 is strongly dependent on both the edited site and cellular context. Together, these findings define stoichiometry- and locus-dependent m6A regulation as a general mechanism for tuning gene expression during developmental metabolic transitions in the heart and, likely, other tissues.

## Results

### High-Resolution Nanopore Direct RNA Sequencing Unveils a Bimodal m6A Regulatory Code Governing Gene Expression Stoichiometry in Embryonic Hearts

The precise molecular mechanisms by which epitranscriptomic modifications translate into diverse functional outcomes, particularly during rapid developmental transitions, are largely unknown due to inherent limitations of conventional methodologies^13–15^. To overcome these barriers and achieve an unprecedented level of resolution, we pioneered the application of PromethION high-depth nanopore direct RNA sequencing (DRS) to E12.5 embryonic mouse hearts. This state-of-the-art platform allowed for the direct, single-molecule characterization of m6A sites and, critically, their quantitative stoichiometric ratios across full-length transcripts, bypassing the biases of antibody-based enrichment and RNA fragmentation inherent in previous m6A-Seq techniques^13–15,31^. Our initial analysis confirmed a known transcriptome-wide distribution of m6A sites in control hearts^8,12^, predominantly localized to the 3’UTR (49.8%) and coding sequences (CDS) (38.8%) (Fig. 1A). To establish a robust *in vivo* model for dissecting m6A function, we generated a conditional knockout of the m6A writer Mettl14 in cardiac progenitor cells. Thus, we conditionally deleted the *Mettl14* writer in cardiac progenitor cells *in vivo* using *Nkx2.5-Cre*^32^ mice that do not exhibit a hypomorphic phenotype, to generate *Nkx2.5-Cre*^32^*; Mettl14 fl/fl* (M14-KO) mice (Fig. 1B). The subsequent Mettl14 downregulation (Fig. 1B) resulted in a profound and global reduction in cardiac m6A mRNA levels (Fig. 1C-E, Supplemental Fig. S1), confirming the efficient perturbation of the m6A epitranscriptome.

**Figure 1.**
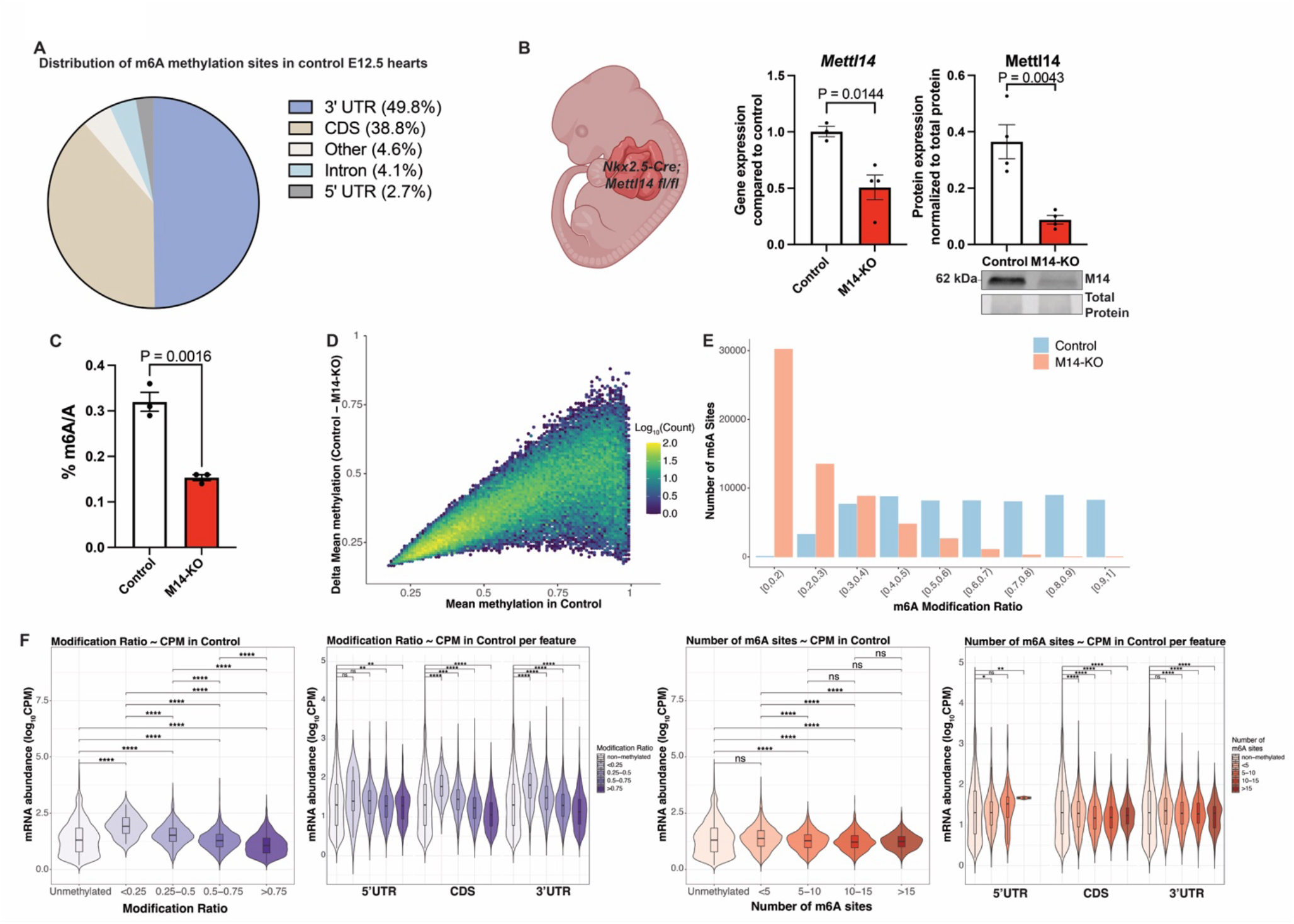
Global m6A landscape captured using Nanopore long-read direct RNA sequencing in E12.5 control and M14-KO hearts. (A) Transcriptome-wide distribution of m6A sites across mRNA features in four control samples, showing predominant enrichment in the 3′ untranslated regions (3′UTRs) and coding sequences (CDS). (B) Schematic illustration of METTL14 conditional knockout in cardiac progenitor cells using Nkx2.5-Cre (M14-KO). Quantification of *Mettl14* transcript (n=3 control and 4 M14-KO) and protein levels (n=4) in control vs M14-KO hearts at E15.5, confirming efficient M14 knockout. (C) Mass-Spectrometry quantification of global m6A/A ratio using E15.5 hearts showing marked loss of m6A methylation in M14-KO (n=3/group). (D) MA plot showing methylation difference (Control minus M14-KO, n=4) as a function of baseline methylation level in control hearts. Data are binned in two dimensions; point color represents log scaled count. Mettl14 depletion leads to a global reduction in m6A mRNA. (E) Bar graphs showing the number of m6A sites across corresponding modification frequency categories. Sites with m6A probability ≥0.9 (calculated from modkit pileup) classified as methylated. Only methylation sites with ≥10 reads in ≥3 biological replicates per condition are represented. A global loss of higher methylation ratios was observed with a distribution shift towards very low m6A modified transcript ratios in M14-KO hearts. (F) Violin plots showing mRNA abundance (CPM) of transcripts grouped by their modification ratio or number of m6A sites per gene at baseline. Both were further stratified by the subcellular location of the m6A site (5′UTR, CDS, or 3′UTR). Transcripts with lower modification ratios (<0.5) showed higher abundance compared to non-methylated mRNAs suggesting m6A-mediated mRNA upregulation. In contrast, higher m6A ratios (>0.5) or more than 5 m6A sites per transcript were associated with reduced abundance consistent with m6A-mediated transcript decay. This trend was consistent for m6A sites in CDS and 3’UTR regions. 5’UTR regions had an overall lower number of m6A sites, and the ones with higher m6A counts were associated with higher expression levels. Two-sided Welch’s t-tests, with multiple-testing correction by Benjamini–Hochberg (BH) were performed. ∗p < 0.05, ∗∗p < 0.01, ∗∗∗p < 0.001, and ∗∗∗∗p < 0.0001. For B and C, all quantitative data are presented as mean ± SEM. The Shapiro-Wilk test was performed to assess normal distribution, and student parametric t-test (two-tailed) or Mann-Whitney (two-tailed) non-parametric tests were used as appropriate for all comparisons. Only *P* values <0.1 are reported. For E and F, GO enrichment adjusted *P* values were corrected using BH to control false discovery rate. Gene ratio is defined as the number of genes in the differentially expressed list divided by the total number of genes annotated to the respective GO term.

m6A epitranscriptomics can regulate transcript levels, thus we comprehensively assessed how m6A transcript ratios affect mRNA abundance (Fig. 1F). We averaged the modification ratio of all identified m6A sites per transcript and formed five subgroups of genes (unmethylated, <0.25, 0.25-0.5, 0.5-0.75 and >0.75). This comprehensive quantitative nanopore DRS analysis, uniquely enabled by this high-resolution approach, revealed a novel bimodal regulatory code for m6A, where distinct modification ratios dictate divergent molecular outcomes on gene expression. We found that transcripts bearing low m6A modification ratios (<0.5) were consistently associated with significantly increased mRNA abundance, suggesting a novel m6A-mediated mechanism for enhancing transcript stability or translation. Conversely, transcripts with higher m6A ratios (>0.5) were tightly linked with lower transcript levels, indicative of canonical m6A-mediated suppression or decay (Fig. 1F). This bimodal effect was robustly observed across various genomic features, being particularly pronounced for transcripts harboring m6A sites within CDS and 3’UTR (Fig. 1F). Extending this finding, we also categorized m6A modified mRNAs in five groups depending on the number of methylation sites per transcript (unmethylated, <5, 5-10, 10-15, >15) and discovered that transcripts with a higher density of m6A sites (>5 per transcript) generally correlated with reduced mRNA abundance, predominantly at 3’UTR sites. Intriguingly, a limited subset of 5’UTR m6A-dense transcripts exhibited an opposite, positive correlation with mRNA abundance (Fig. 1F). These quantitative insights, directly challenging the prevailing view of a universal m6A effect, establish a refined paradigm for epitranscriptomic regulation, highlighting the critical, context-dependent role of m6A stoichiometry in shaping mRNA fate and ultimately gene expression.

### m6A Epitranscriptomics is Essential for Cardiac Development

Having uncovered a fundamental bimodal m6A regulatory code, we next sought to determine its physiological significance during critical developmental transitions in the heart. Loss of Mettl14-mediated m6A in cardiac progenitor cells resulted in severe ventricular non-compaction cardiomyopathy, characterized by pronounced trabeculations, thinning of the compact layer, and overall disorganized ventricular morphology (Fig. 2A). These structural anomalies were accompanied by significant functional impairment, with reduced left ventricular systolic function and mild cardiomegaly observed via *in utero* echocardiography at E15.5 (Fig. 2B, Supplemental Fig. S2A). Notably, M14-KO embryos exhibited embryonic lethality by late gestation (Fig. 2C), underscoring the indispensable and non-redundant role of m6A epitranscriptomics in early heart formation. At the cellular level, M14-KO hearts displayed significantly impaired cardiomyocyte proliferation shown by reduced phospho-histone 3 (pH3, mitosis marker) and EdU (S phase marker) cells (Fig. 2E) and a concomitant increase in apoptosis (TUNEL+ cells) (Fig. 2F), likely contributing to the observed developmental defects. To further facilitate mechanistic dissection, we developed a robust *in vitro* model of Mettl14 knockdown (M14-KD) in hiPSC-CMs. Achieved via adenoviral siRNA delivery (Fig. 2F), this led to a significant reduction of global m6A mRNA levels (Fig. 2G), accurately recapitulating the m6A depletion observed *in vivo* in M14-KO hearts (Fig. 1C). Importantly, these M14-KD hiPSC-CMs also mirrored the *in vivo* cellular phenotypes, displaying significantly reduced proliferation (Fig. 2H). Thus, our comprehensive *in vivo* and *in vitro* data conclusively demonstrate that m6A epitranscriptomics is a critical and non-redundant regulator, essential for the fundamental processes underpinning normal cardiac development.

**Figure 2.**
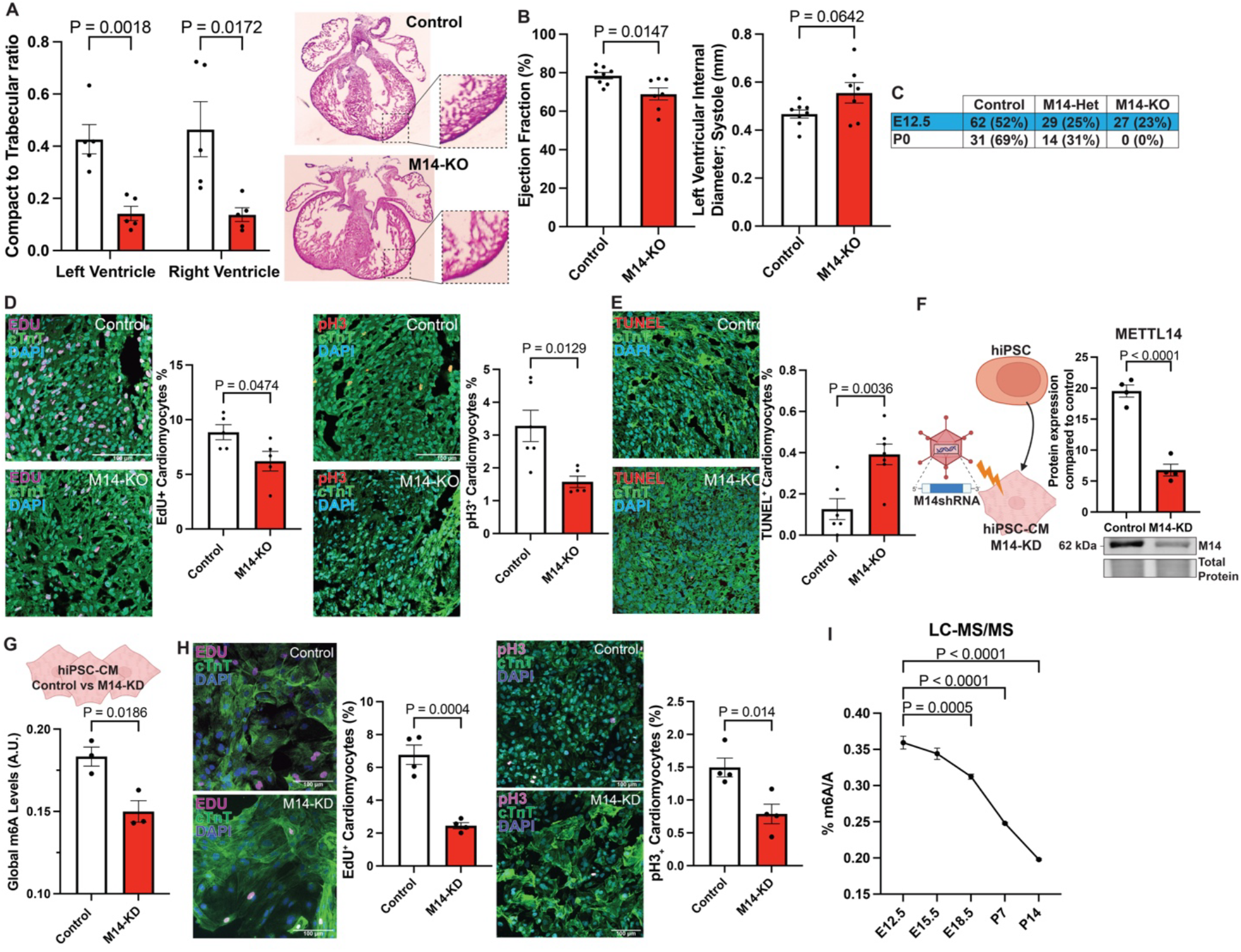
m6A epitranscriptomics is essential for fundamental processes underpinning normal cardiac development. (A) Representative hematoxylin and eosin (H&E)–stained sections and quantification (n=5) of myocardial wall thickness at E15.5 showing reduced compact to trabecular ratio consistent with ventricular non-compaction in M14-KO hearts. (B) In utero echocardiography of E15.5 hearts showing decreased ejection fraction (EF) and altered left ventricular internal diameter at systole (LVID;s) in M14-KO embryos, indicating compromised cardiac contractile function (n=8 control, 7 M14-KO). (C) The table summarizes predicted and observed genotypes at E12.5 and P0, revealing perinatal lethality in M14-KO pups. (D-E) Immunofluorescence staining for phospho-histone H3 (pH3, mitosis), EdU (S-phase), and TUNEL (apoptosis) in control and M14-KO hearts at E15.5 (n ≥ 5). M14-KO hearts showed reduced pH3+ and Edu+ cardiomyocytes consistent with reduced proliferation, while the percentage of TUNEL+ myocytes was increased suggestive of increased apoptosis. cTnT: cardiac Troponin T, DAPI: nuclei. Scale bars: 100 µm. (F) Schematic illustration showing infection of hiPSC-derived cardiomyocytes (CMs) with adenovirus carrying METTL14-shRNA (Ad_M14sh) (M14-KD) or mock adenovirus (Ad_Mock) (control). Western blot quantification confirmed efficient M14 downregulation in M14-KD hiPSC-CMs compared to control. (G) Global m6A quantification showing decreased total m6A levels in M14-KD hiPSC-CMs relative to control (n=3). (H) Representative immunofluorescence images and quantification of proliferative markers in control and M14-KD hiPSC-CMs. Cells were stained for phospho-histone H3 (pH3, Mitosis) or EdU (S-phase), cardiac troponin T (cTnT; cardiomyocyte marker), and DAPI (nuclear stain). Scale bars: 100 µm. Quantification was performed by quantifying pH3- or EdU-positive cardiomyocyte nuclei normalized to total DAPI-positive nuclei (n=4). Reduced proliferation was observed in M14-KD hiPSC-CMs. (I) Mass-Spectrometry quantification of global m6A/A ratio across different embryonic and postnatal murine heart development time points (n=3) showing a progressive decline in m6A methylation during cardiac maturation. All quantitative data are presented as mean ± SEM. The Shapiro-Wilk test was performed to assess normal distribution, and student parametric t-test (two-tailed) or Mann-Whitney (two-tailed) non-parametric tests were used as appropriate for all comparisons. Only *P* values <0.1 are reported.

Supporting this essential role, quantitative analysis using liquid chromatography coupled with mass spectrometry (LC-MS/MS) revealed a dynamic expression profile of global m6A levels during cardiogenesis (Fig. 2I). m6A levels were significantly elevated during early-to-mid embryonic stages (E12.5 and E15.5) before declining postnatally (Fig. 2I). This precisely coordinated regulation, correlating with stable expression of m6A mRNA writer proteins during mid-gestation (Supplemental Fig. S2B), indicated that m6A epitranscriptomics is actively modulated during this critical developmental window, reinforcing the profound impact of its perturbation.

### m6A Epitranscriptomics Critically Orchestrates the Cardiac Metabolic Switch During Development

To unravel the precise molecular pathways underlying these profound developmental and cellular defects, and dissect the spatial and heterogeneous effects of m6A epitranscriptomics, we performed single-cell RNA-Seq using E12.5 mouse hearts^33^, which allowed us to focus on early differentiated cardiomyocytes (CMs) at mid-gestation (Fig. 3A-B, Supplemental Fig. S3A) and circumvent potential secondary effects inherent in later developmental stages. Our single-cell RNA-Seq data revealed that m6A epitranscriptomics critically orchestrates the vital mid-gestation cardiac metabolic switch from glycolysis to OXPHOS^34–38^. Our scRNA-Seq data identified four distinct cardiomyocyte (CM) subpopulations (Fig. 3C). Importantly, upon Mettl14 deletion, we observed a significant alteration in the composition of these CM clusters: the population of CMs characterized by high aerobic respiration (CM2) was markedly decreased in M14-KO hearts (from 44% to 27%), while a less metabolically active CM1 subpopulation expanded (Fig. 3C). Differential gene expression analysis within M14-KO CMs further demonstrated a distinct and detrimental shift in metabolic programming, with a profound downregulation of mitochondrial electron transport, ATP production, and OXPHOS pathways (Fig. 3D-E, Supplemental Fig. S3B). This provided high-resolution evidence for a severe disruption of the metabolic switch at the single-cell transcriptomic level, directly linking m6A epitranscriptomics to the regulation of CM metabolic heterogeneity.

**Figure 3.**
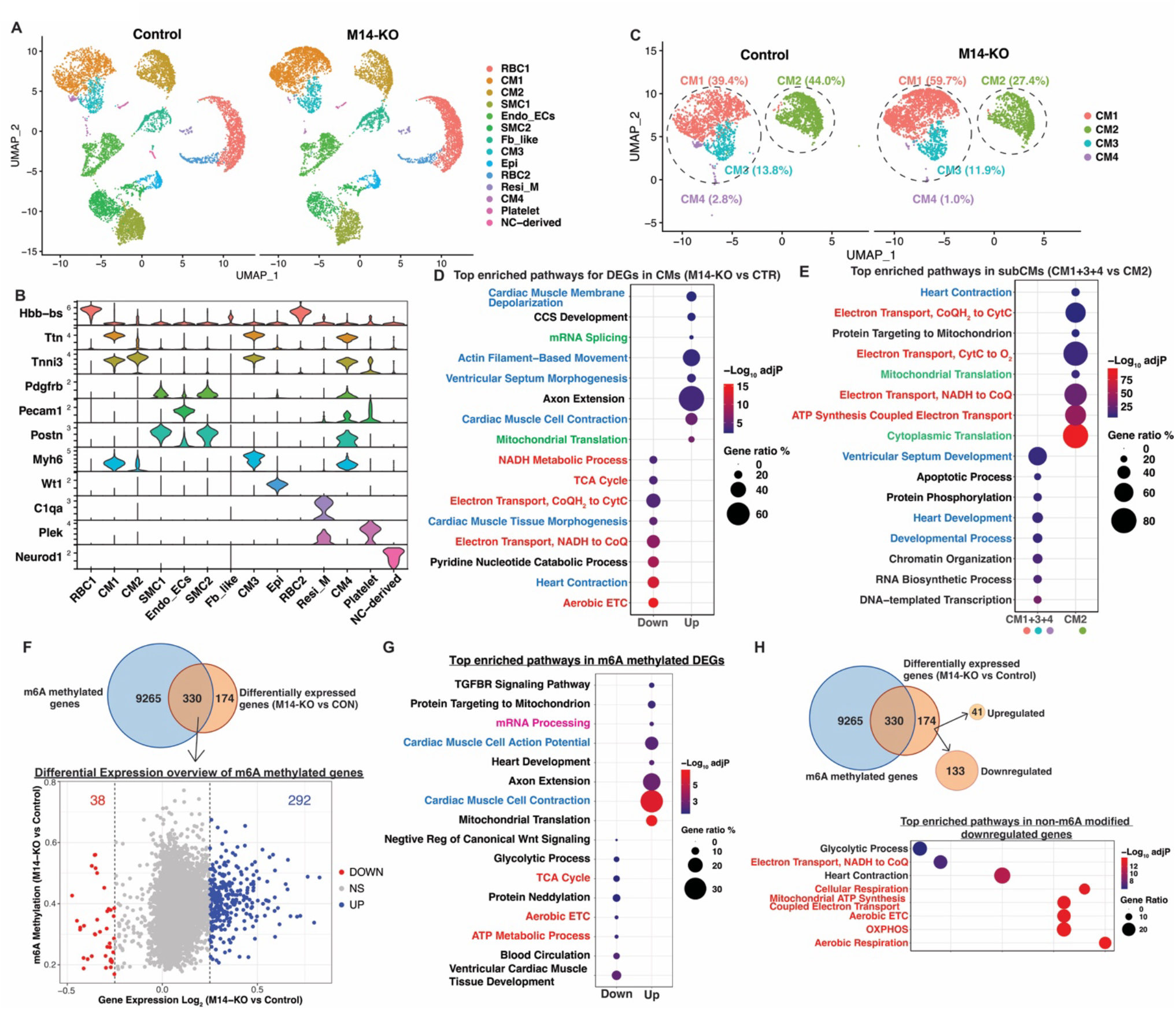
Single-cell transcriptomics of E12.5 M14-KO hearts primarily reveals suppressed aerobic metabolic pathways. (A) UMAP visualization of single-cell transcriptomes from control and M14-KO embryonic hearts at E12.5, colored by major cardiac cell types. RBC: red blood cells; CM: cardiomyocytes; SMC: smooth muscle cells; Endo_ECs: endocardial endothelial cells; Fb: Fibroblasts; Epi: epicardial cells; Resi_M: resident macrophage; NC: neural crest. (B) Violin plots showing expression of representative marker genes used to identify each cell population. (C) UMAP showing cardiomyocyte (CM) subpopulations and quantification of the fraction of each CM cluster in control and M14-KO hearts, indicating altered CM composition upon METTL14 loss. Notably, M14-KO hearts contained a higher percentage of CM1 subpopulation (59.7 vs 39.4%), while CM2 cluster was higher in control hearts (44% vs 27.4%). (D) Differential gene-expression and Gene Ontology (GO) pathway analysis comparing control and M14-KO CMs. M14-KO CMs consistently displayed downregulation of mitochondrial electron-transport and ATP-production pathways (red) and upregulation of gene expression (green). CM contraction pathways (blue) showed a mixed pattern as actin-filament and depolarization pathways were upregulated, whereas cardiac morphogenesis regulation and heart contraction pathways were reduced. (E) GO pathway analysis of the CM2 subpopulation, which was proportionally higher in control hearts, showed enrichment of oxidative metabolism and translation pathways and reduced expression of transcription-related processes. (F) Overlap of m6A-modified and differentially expressed genes in cardiomyocytes suggesting direct m6A-mediated regulation. Volcano plot of methylation change (Control minus M14-KO) versus log2 fold change in gene expression. Genes with |log2FC| ≥0.25 and adjusted p ≤0.05 are shown; vertical dashed lines indicate expression cutoffs. Most m6A differentially expressed genes were up-regulated in M14-KO hearts consistent with m6A-mediated mRNA decay. (G) GO pathway analysis of m6A modified genes that are differentially expressed in M14-KO hearts. Upregulated pathways include cardiac muscle contraction (blue) and mRNA processing (magenta), while the small number of suppressed genes showed enrichment of oxidative metabolism pathways (red). (H) Identification of genes that are differentially expressed in M14-KO hearts but are lacking detectable m6A modification consistent with indirect transcriptional regulation. The majority of the genes were downregulated. GO pathway analysis of non–m6A methylated downregulated genes in M14-KO hearts shows enrichment of aerobic metabolism pathways (red). *P*-values were adjusted by Benjamini-Hochberg method in (D, E) and gene ratio as the number of genes in the differentially expressed list divided by the total number of genes annotated to the respective GO term.

To dissect the mechanisms of m6A-mediated gene regulation, we overlapped differentially expressed genes with m6A-methylated transcripts from our nanopore DRS data (Fig. 3F). This analysis revealed that most of the upregulated genes in M14-KO CMs were m6A-modified, while only a minor proportion of downregulated transcripts overlapped with m6A methylation. This finding suggests that, in this developmental context, m6A primarily drives mRNA decay (Fig. 3F). Pathway analysis of these m6A-modified genes highlighted significant metabolic changes, specifically revealing enhanced mitochondrial protein synthesis and increased expression of cardiac muscle genes. Concurrently, there was a pronounced repression of genes for the electron transport chain (ETC), TCA cycle, and glycolysis, as well as those regulating ventricular muscle development (Fig. 3G). Furthermore, examination of suppressed, non-m6A-modified genes indicated their predominant involvement in aerobic respiration, with a central impact on the ETC (Fig. 3H). This indicates that the m6A epitranscriptome directly and indirectly regulates aerobic metabolism gene expression during embryonic heart development.

A substantial number of genes possess m6A modifications that do not manifest as changes in transcript level (Fig. 3F); thus, complementing these transcriptomic insights, we utilized mass spectrometry proteomics on E12.5 M14-KO hearts to comprehensively assess protein-level changes and potential translational regulation (Fig. 4A). Proteomics revealed a clear signature of reduced aerobic metabolism, with specific and widespread downregulation of major subunits across all mitochondrial ETC complexes, particularly Complex I and II (Fig. 4B-C). Importantly, our integrated analyses revealed a significant and widespread discordance between mRNA and protein abundance for numerous genes in M14-KO hearts (Fig. 4D), strongly implicating m6A in translational and other post-transcriptional regulatory mechanisms. For example, several sarcomeric genes showed altered mRNA levels but no significant change in corresponding protein levels, pointing towards mRNA-level-only regulation (Fig. 4D, Supplemental Fig. S4A-B). Pathway enrichment analysis of m6A-methylated genes with protein-level-only differential expression further highlighted downregulation of metabolism and protein processing (Fig. 4E). These protein-level findings powerfully corroborated the metabolic disruption observed at the transcriptomic level.

**Figure 4.**
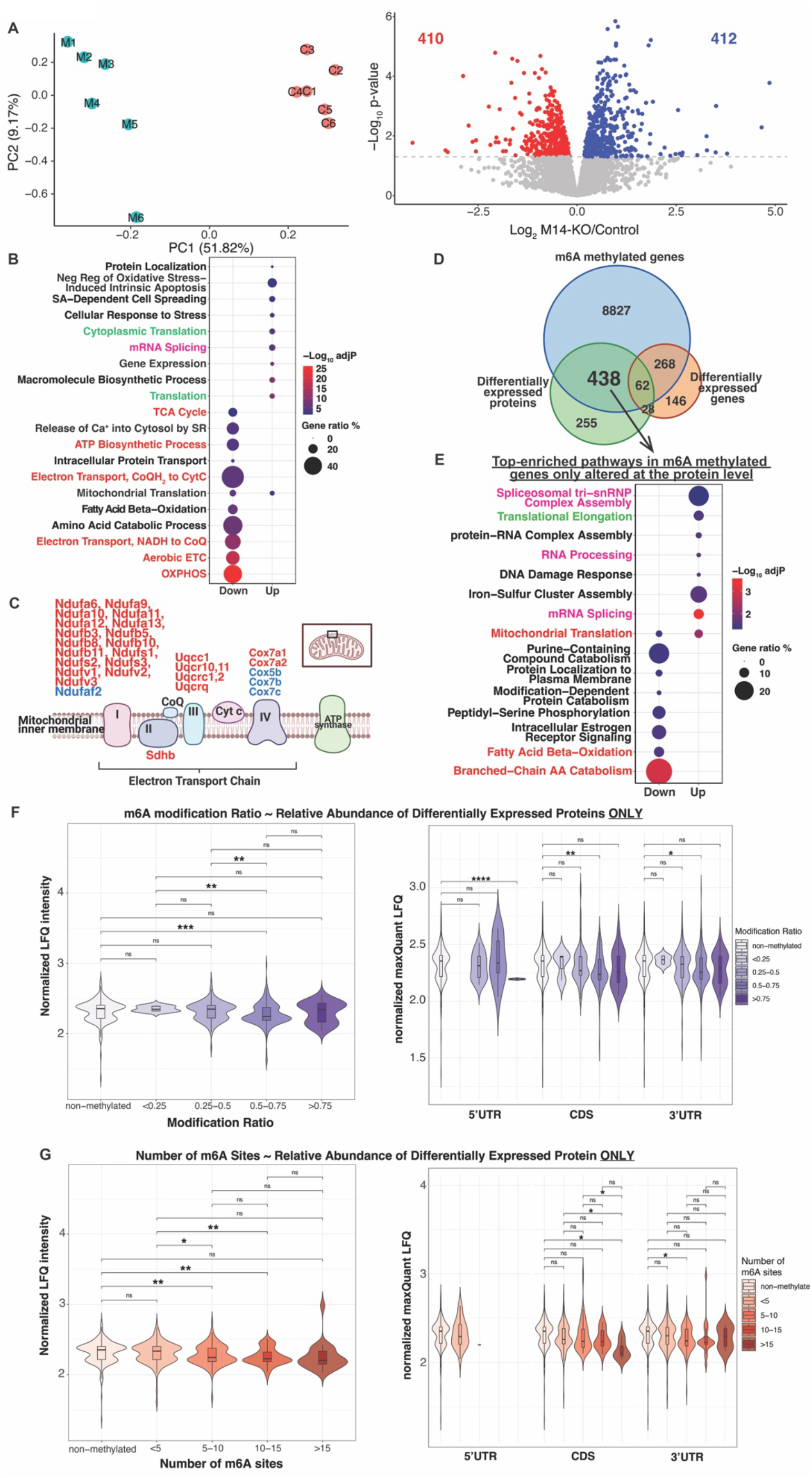
Proteomics analyses of M14-KO vs control E12.5 hearts uncovers m6A epitranscriptomics regulation at the protein level. (A) Principal component analysis (PCA) of proteomic profiles from control and M14-KO E12.5 hearts (n=6), showing clear separation between groups. Volcano plot of differentially expressed proteins. The -log10 p values are derived from empirical Bayes–moderated p values (p.mod) generated by limma (moderated t statistic using variance shrinkage and increased degrees of freedom). (B) GO pathway analysis of differentially expressed proteins, highlighting upregulated translation (green) and mRNA processing (magenta) pathways and downregulated aerobic metabolism pathways (red). Mitochondrial electron transport chain (ETC) proteins were primarily decreased. (C) Illustration of mitochondrial ETC complexes, listing the downregulated (red) and upregulated (blue) ETC components. Complex I subunits were the predominate downregulated ETC proteins. (D) Overlap of genes and proteins identified across three datasets: m6A-modified transcripts (nanopore; m6A probability ≥0.9), differentially expressed transcripts (scRNA-seq; |log2FC| ≥0.25 and adjusted p ≤0.05), and differentially expressed proteins (mass spectrometry; |log2FC| ≥0.25 and p.mod ≤0.05). There is minimal overlap between the set of differentially expressed proteins and the m6A-modified, differentially expressed genes. The observed changes in protein expression for the majority of differentially expressed proteins occurred without significant alterations in their corresponding m6A-modified transcript levels, indicating a mechanism of m6A-mediated translational control for these proteins. (E) Functional enrichment analysis (GO pathways) of m6A-methylated genes, which displayed protein-level-only differential expression indicative of m6A-mediated translational control, showed significant downregulation of pathways associated with metabolism and protein processing (red). Conversely, upregulated proteins were predominantly enriched in functions related to translation (green) and mRNA processing (magenta). (F-G) Violin plots of normalized protein abundance (LFQ intensity) grouped by m6A modification ratio or number of m6A sites per transcripts, further clustered based on m6A site location (5′UTR, CDS, 3′UTR). Higher ratios (>0.5) and number of m6A sites (>5) were consistent with lower protein abundance. Statistical analyses using Two-sided Welch’s t-test with Benjamini–Hochberg (BH) correction for multiple comparisons were performed. Significance is denoted as: ∗p < 0.05, ∗∗p < 0.01, ∗∗∗p < 0.001, ∗∗∗∗p < 0.0001.

To distinguish m6A-mediated translational regulation from changes in mRNA expression for specific targets, we meticulously correlated the abundances of differentially expressed proteins observed in M14-KO hearts with m6A transcript motifs, focusing on those proteins whose corresponding mRNA levels remained unchanged (Fig. 4D). This analysis provided deep insight into the functional consequences of m6A patterning on translation. We found a clear inverse correlation between normalized protein abundance and m6A modification ratios: specifically, higher m6A ratios (0.5–0.75) within the coding sequence and 3’ UTR were associated with decreased protein levels (Fig. 4F). Similarly, transcripts harboring a greater number of m6A modifications (5-15 per transcript) in their CDS and 3’UTR regions consistently demonstrated lower levels of protein expression (Fig. 4G). Notably, this pattern was not observed for 5’UTR m6A sites. These findings strongly indicate that specific m6A modification patterns exert distinct effects by inhibiting protein synthesis of target transcripts, serving as a critical post-transcriptional control mechanism.

To functionally validate these multi-omics findings and gain a deeper mechanistic understanding of metabolic impairment in CMs, we performed comprehensive metabolic profiling in our M14-KD hiPSC-CMs (Fig. 5A). Using Seahorse XFe96 Analyzer, M14-KD hiPSC-CMs displayed significantly compromised OXPHOS capacity, evidenced by lower basal and maximal oxygen consumption rates (OCR) compared to controls (Fig. 5B). This impairment was specific to aerobic respiration, as extracellular acidification rate (ECAR), a measure of glycolysis, remained unchanged (Fig. 5C), aligning with our *in vivo* omics data. Further resolution using substrate utilization assays in permeabilized hiPSC-CMs revealed impaired function of specific ETC complexes I and II, but not Complex III or IV, as indicated by reduced ADP-stimulated OCR and Respiratory Control Ratio (RCR) (Fig. 5D-E). No difference in mitochondria content was detected in either M14-KD hiPSC-CMs or M14-KO E12.5 hearts (Supplemental Fig. S5A-B). This detailed functional validation in hiPSC-CMs powerfully corroborates our *in vivo* findings and establishes a direct and essential link between m6A epitranscriptomics and the precise molecular control of cardiac metabolic maturation, a prerequisite for healthy heart development.

**Figure 5.**
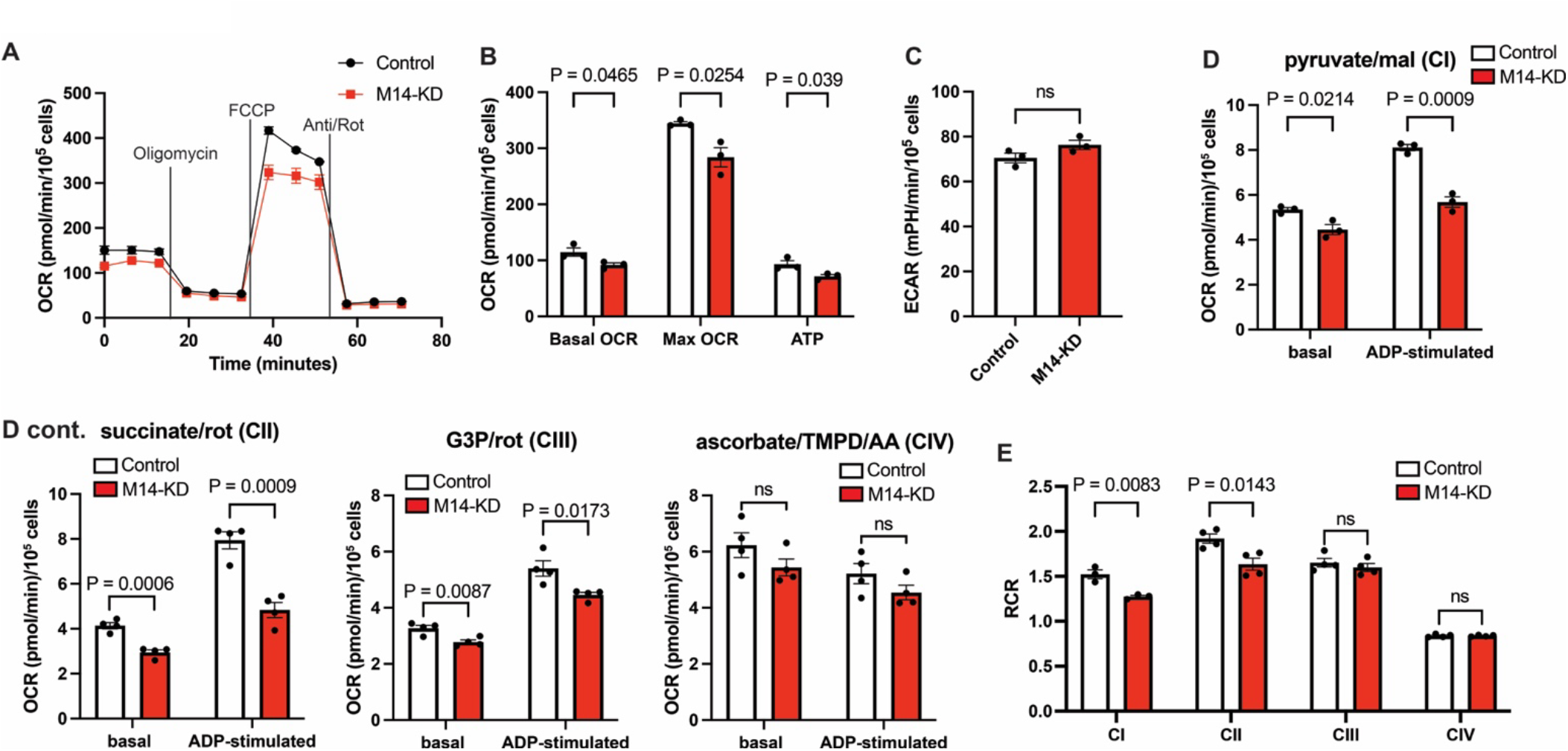
M14-KD in hiPSC-CMs results in decreased oxidative phosphorylation. (A) Electron flow through the respiratory chain in intact mitochondria. Injections of OXPHOS-targeting agents and metabolic modulators were performed as indicated. FCCP: Carbonyl cyanide-4-(trifluoromethoxy)phenylhydrazone; Oligo: oligomycin, AA: Anti: antimycin A; Rot: rotenone. (B) Quantification of metabolic parameters including basal oxygen consumption rate (OCR), maximum OCR and ATP production (n=3). All parameters were reduced in M14-KD hiPSC-CMs indicating impaired oxidative metabolism. (C) Extracellular acidification rate (ECAR) measurements (n=3) showed no significant change in M14-KD hiPSC-CMs, confirming that glycolysis was not affected (D-E) Quantification of basal and ADP-stimulated respiration (G) and respiratory control ratio (RCR) (H) in permeabilized CMs using the specified substrates and rotenone as listed (n ≥ 5 wells from at least 3 independent experiments) showed that M14 depletion in hiPSC-CMs results in impaired function of ETC complexes I, II, and III, while complex IV activity remains intact, highlighting a selective disruption of mitochondrial respiration. All quantitative data are presented as mean ± SEM. The Shapiro-Wilk test was performed to assess normal distribution, and student parametric t-test (two-tailed) or Mann-Whitney (two-tailed) non-parametric tests were used as appropriate for all comparisons. Only *P* values <0.1 are reported.

### m6A Epitranscriptomics-YY1 Regulatory Axis Post-Transcriptionally Drives Metabolic Reprogramming

With the profound metabolic disruption resulting from Mettl14-mediated m6A loss firmly established, we next sought to identify the specific molecular effector linking m6A depletion to this phenotype. Our integrated multi-omics analysis (transcriptomics and proteomics) revealed a significant and widespread discordance between mRNA and protein abundance for numerous genes in M14-KO hearts (Fig. 4D), strongly implicating m6A in translational or other post-transcriptional regulatory mechanisms. Crucially, we observed that a substantial portion of the downregulated OXPHOS genes in M14-KO hearts did not themselves harbor m6A modifications (Fig. 3H). This key finding prompted our hypothesis that m6A epitranscriptomics might exert its effect indirectly by regulating key transcription factors (TFs) that, in turn, control OXPHOS gene expression.

To identify such TFs, we performed a systematic computational analysis. We integrated our scRNA-Seq data (Fig. 3) with chromatin immunoprecipitation (ChIP) data from ENCODE^39^ and ChEA^40^ projects, specifically looking for TFs enriched in the promoters of the non-m6A modified suppressed genes (Fig. 6A). Among the compelling candidates identified (e.g., Brca1, Pml), Yin Yang 1 (YY1) emerged as a particularly strong candidate. It can function as a pivotal transcriptional repressor with well-established roles in cellular metabolism^41–46^, it was present in our m6A-modified transcript pool, and crucially, exhibited high expression specifically in CMs (Fig. 6B). Further reinforcing this, our scRNA-Seq analysis revealed that E12.5 CMs from metabolically active cluster 2, which displayed increased OXPHOS gene expression, exhibited exceptionally low levels of YY1 (Supplemental Fig. S6A), suggesting an inverse correlation with metabolic maturation. This convergence of evidence, coupled with previous studies demonstrating a decline in YY1 during heart development^17^ (Supplemental Fig. S6B) and sustained high YY1 levels in immature hiPSC-CMs^29^ (Supplemental Fig. S6C), strongly supported our hypothesis that elevated YY1 can function as a repressor of OXPHOS genes in this developmental setting.

**Figure 6.**
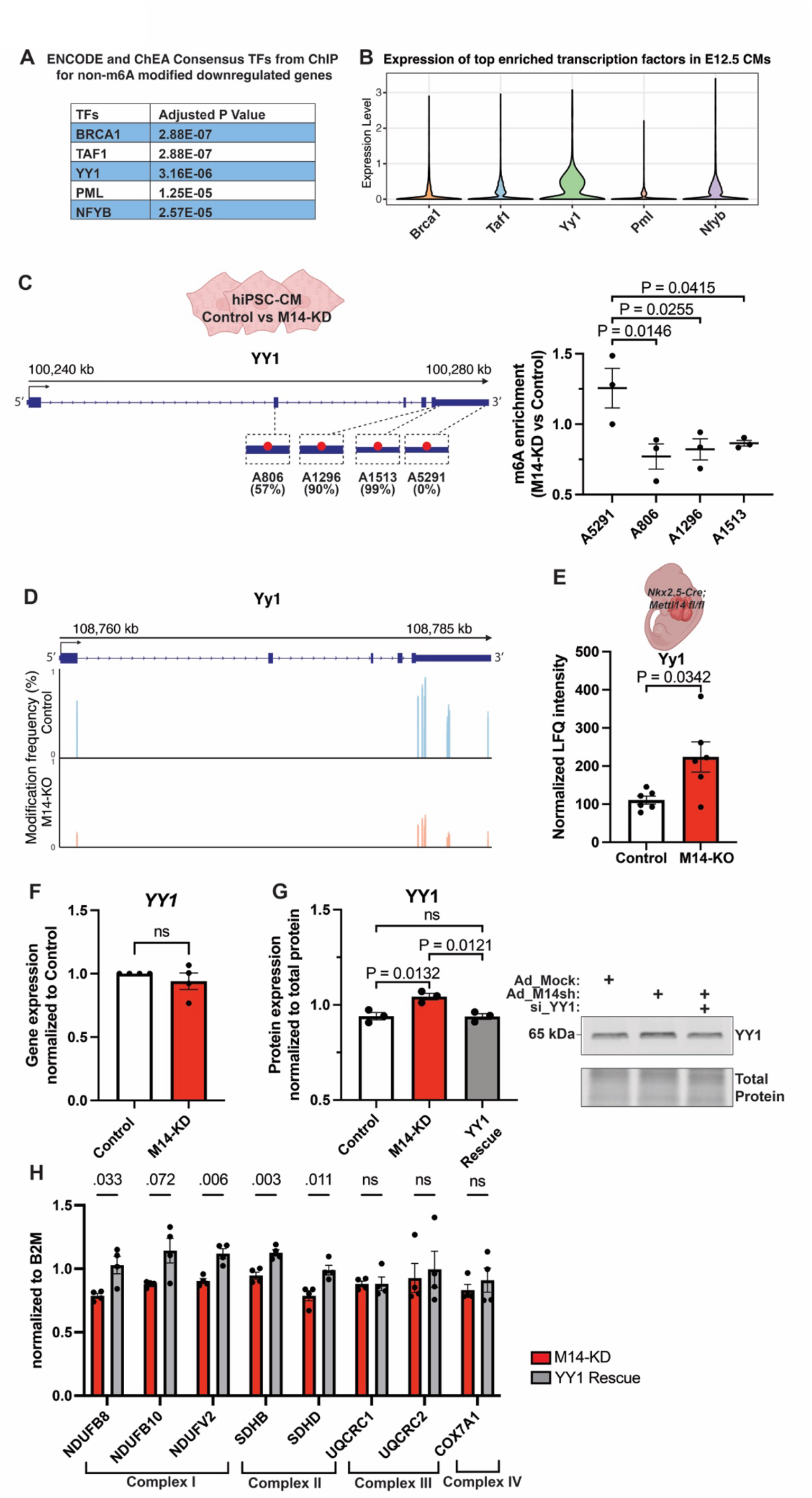
>m6A epitranscriptomics regulates YY1 expression, linking m6A to mitochondrial oxidative metabolism during cardiac development. (A) ENCODE/ChEA transcription factor enrichment for non-m6A methylated downregulated genes. Top enriched TFs include BRCA1, TAF1, YY1, PML, and NFYB (adjusted p values shown in accompanying table). (B) Among the five most enriched transcription factors, *Yy1* exhibited highest expression levels in E12.5 cardiomyocytes *in vivo* based on single cell RNAseq. (C) m6A RIP–qPCR analysis of *YY1* mRNA in M14-KD hiPSC-CMs. Four primer pairs were designed: three targeting exonic or 3’ UTR regions containing predicted m6A sites (DeepSRAMP predictions, probability values indicated in %) and one targeting a non-m6A region. M14 knockdown resulted in reduced m6A enrichment at all three predicted sites, whereas no change was observed at the non-m6A site (n=3). (D) m6A methylation profiles of *Yy1* in E12.5 control vs M14-KO mouse hearts based on nanopore m6A-Seq showing m6A methylation peaks at 5’ and 3’ UTR regions. All high probability m6A sites displayed profound decrease in M14-KO hearts. (E) Quantitative proteomics showed a significant increase in Yy1 protein level in M14-KO E12.5 hearts compared to controls. (F) *YY1* gene expression remained unchanged in M14-KD hiPSC-CMs (n=4). (G) Sequential knockdown experiment used to assess the role of YY1 in ETC gene regulation. hiPSC-CMs were first transduced with shRNA-carrying adenovirus to knock down *METTL14*, followed by siRNA transfection of either siNC (non-targeting negative control) or siYY1 (targeting YY1) 12 hours later. Western blot quantification showed increased YY1 protein levels in M14-KD hiPSC-CMs + siNC compared to control + siNC and M14-KD + siYY1 cells (n=3). (H) Gene expression analyses in hiPSC-CMs revealed that the suppression of ETC complex I and II genes in M14-KD hiPSC-CMs was partially alleviated by subsequent *YY1* knockdown (n=3). These findings suggest that the m6A-mediated regulation of YY1 plays a key role in controlling the expression of ETC genes likely accounting for the decreased oxidative metabolism observed in M14-KD hiPSC-CMs. All quantitative data are presented as mean ± SEM. The Shapiro-Wilk test was performed to assess normal distribution, and student parametric t-test (two-tailed) or Mann-Whitney (two-tailed) non-parametric tests were used as appropriate for all comparisons. Only *P* values <0.1 are reported.

To experimentally validate YY1 as an m6A-modified target, we first utilized the deepSRAMP^47^ online prediction platform to identify putative *YY1* m6A methylation sites, leveraging its high-accuracy deep learning algorithm. Subsequently, we performed m6A RNA immunoprecipitation (RIP) in control and M14-KD hiPSC-CMs to precisely quantify m6A enrichment. Our RIP analysis identified three *YY1* m6A modification sites (two within the coding region and one in the 3’UTR) that were significantly enriched in control hiPSC-CMs and profoundly decreased in M14-KD cells (Fig. 6C), directly confirming the presence of Mettl14-dependent m6A modification on the *YY1* transcript. Our nanopore DRS analysis of M14-KO hearts comprehensively corroborated these findings, confirming a profound decrease in m6A at these same *Yy1* sites within both coding and 3’ UTRs (Fig. 6D). Critically, this loss of m6A on *Yy1* transcripts directly correlated with a substantial increase in Yy1 protein levels in M14-KO hearts (Fig. 6E). Importantly, *Yy1* transcript levels remained unchanged (Fig. 6F, Supplemental Fig. S6D), and other identified transcription factors showed no significant effects (data not shown), strongly indicating that m6A epitranscriptomic regulation specifically and post-transcriptionally modulates YY1 protein expression. This inverse correlation between *YY1* transcript m6A and YY1 protein levels aligns with our newly defined bimodal m6A code, where higher m6A ratios in CDS/3’UTR consistently correlated with lower protein abundance (Fig. 4F, 4G). This collectively establishes a novel m6A-YY1 regulatory axis, where m6A epitranscriptomics precisely controls the activity of a key transcriptional repressor.

To functionally confirm the causality of this m6A-YY1 axis in driving metabolic reprogramming and to investigate the direct link between m6A epitranscriptomics and ETC gene regulation via YY1, we performed siRNA-mediated knockdown experiments to normalize the elevated YY1 expression in M14-KD hiPSC-CMs. To mitigate potential secondary effects from excessive YY1 reduction, we first confirmed that the 24-hour siRNA treatment resulted in YY1 protein levels comparable to baseline (Fig. 6G). This precise normalization successfully rescued the expression of representative ETC Complex I and II genes (Fig. 6H). This finding confirms that m6A-mediated regulation of YY1, acting as a repressor, plays a critical role in controlling ETC gene expression and thereby facilitating the crucial aerobic metabolic transition during cardiogenesis.

### A Pioneering hiPSC-Based Platform Reveals Context-Dependent Effects of Targeted m6A Regulation on YY1 Expression

Finally, to directly validate the m6A-YY1 axis and investigate the intrinsic context-dependency of m6A function with unparalleled precision, we developed and deployed a novel CRISPR-based hiPSC platform for targeted epitranscriptomic editing. This platform involved engineering hiPSCs to stably express a dCas13-Mettl3 (dCas13-M3) fusion system, enabling highly specific m6A installation at desired transcript sites (Fig. 7A). We first demonstrated the platform’s efficacy by showing that targeted m6A methylation of the previously identified A1216 m6A methylation site within the beta-actin (*ACTB*) transcript^48^ 3’UTR, efficiently reduced *ACTB* mRNA levels in hiPSC-CMs, consistent with m6A-mediated mRNA degradation mechanisms (Fig. 7B). Moving to our core finding, by precisely targeting specific m6A sites within the *YY1* transcript (identified via RIP-qPCR, Fig. 7C) using our dCas13-M3 platform, we observed a direct downregulation of YY1 protein expression in hiPSC-CMs (Fig. 7C). This result mirrored our *in vivo* M14-KO and *in vitro* M14-KD findings, serving as direct validation of the m6A-YY1 axis. Importantly, these effects were observed exclusively at the protein level (Fig. 7D), further substantiating the post-transcriptional nature of m6A’s control over YY1.

**Figure 7.**
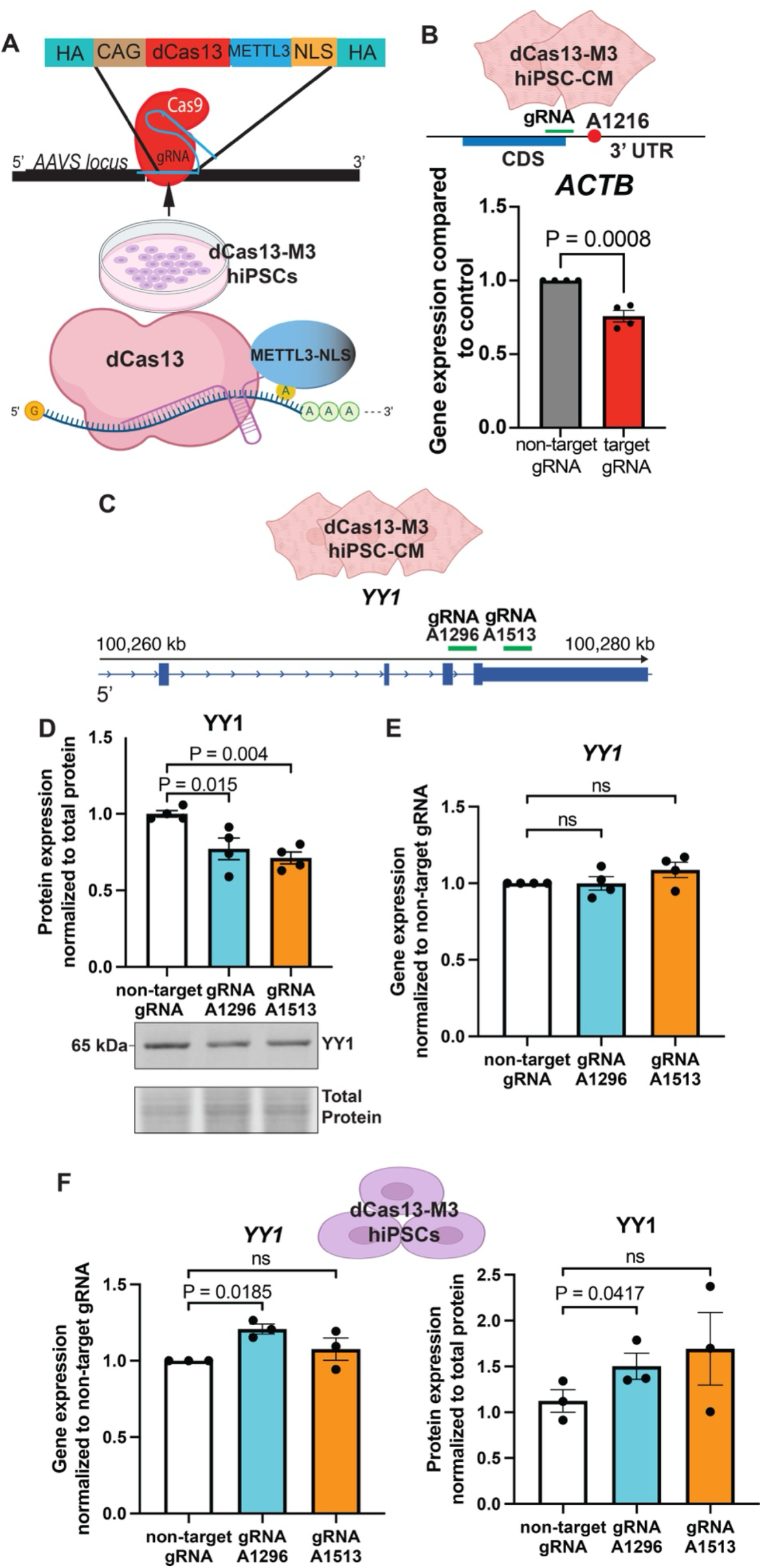
A novel hiPSC-Based Platform for targeted epitranscriptomic editing reveals context-dependent m6A regulation on YY1. (A) CRISPR/Cas9 was used for knock-in of CAG promoter with dCas13-Mettl3 and nuclear localizing sequence (NLS) into the safe *AAVS* locus to prevent silencing. (B) gRNAs targeting the *ACTB* 3’UTR region or control were used. qPCR showed down-regulation of the *ACTB* consistent with m6A-mediated mRNA decay (n=4/group). (C) Schematic representation of gRNAs targeting previously identified *YY1* mRNA m6A methylation sites for targeted epitranscriptomic editing in dCas13-M3 hiPSC-CMs. (D) Targeted m6A epitranscriptomics editing of *YY1* in dCas13-M3 hiPSC-CMs using specific gRNAs for the coding (A1296) and 3’UTR (A1513) resulted in protein downregulation (n=4). (E) *YY1* gene expression remained unchanged in hiPSC-CMs dCas13-M3 after targeted epitranscriptomic editing with gRNAs for CDS (A1296) and 3’ UTR (A1513) (n=4). (F) YY1 gene and protein expression in dCas13-M3 hiPSCs post epitranscriptomic editing, showing increased YY1 gene and protein levels only when targeting the exonic A1296 site. (n=3) All quantitative data are presented as mean ± SEM. The Shapiro-Wilk test was performed to assess normal distribution, and student parametric t-test (two-tailed), Mann-Whitney (two-tailed) non-parametric tests or one-way ANOVA with Bonferroni’s multiple comparisons test were used as appropriate for all comparisons. For single cell RNA-Seq analysis statistics were performed as described in Methods. Only *P* values <0.1 are reported.

The regulatory outcome of targeted m6A editing was found to be exquisitely sensitive to both the precise editing site location within the *YY1* transcript and the cellular context. While targeted m6A methylation consistently suppressed YY1 protein in differentiated hiPSC-CMs (Fig. 7C), applying the same editing strategies to undifferentiated hiPSCs did not result in a similar increase in gene and protein expression; instead, it elicited context-dependent differential effects based on the targeted region (Fig. 7E). This striking divergence provides compelling evidence that m6A-mediated gene regulation is fundamentally shaped by its specific genomic locus and the prevailing developmental stage or cellular state. It robustly demonstrates that m6A signals are dynamically interpreted by the cellular context, yielding adaptive and nuanced regulatory outcomes, rather than a fixed, universal response. This innovative hiPSC-based platform, therefore, represents an invaluable, high-resolution resource for dissecting and manipulating complex, context-dependent RNA-based regulatory mechanisms across diverse hiPSC-derived tissues, offering unprecedented control and clarity for future studies aimed at unraveling epitranscriptomic codes in human development and disease.

## Discussion

The remarkable precision of gene expression in orchestrating complex biological processes, from embryonic development to metabolic homeostasis, relies on multiple layers of control. While m6A epitranscriptomics is increasingly recognized as a key regulatory mechanism^5–7^, the fundamental rules governing its diverse effects – how site-specificity, stoichiometry, and cellular context dynamically shape mRNA fate – have remained mechanistically elusive. Our study addresses these critical questions by combining pioneering high-resolution nanopore DRS with a novel CRISPR-based hiPSC platform. This comprehensive approach allowed us to uncover a bimodal m6A regulatory code driven by stoichiometry, identify a post-transcriptional m6A-YY1 axis crucial for metabolic transitions, and establish that m6A regulation is exquisitely context-dependent, yielding adaptive molecular outcomes that are essential for cardiac development.

Our discovery of a bimodal m6A regulatory code governed by stoichiometry fundamentally reframes how m6A signals are interpreted. Prior models of m6A function often suggested a dominant role in mRNA decay^49^, or a more generalized effect on translation^8–10,50^, frequently based on bulk m6A levels. By leveraging high-depth nanopore DRS, which offers unprecedented single-molecule resolution and quantitative stoichiometry^13–15^, our work moves beyond these limitations. We found that distinct modification ratios (low vs. high) on specific transcripts elicit divergent outcomes (enhanced vs. suppressed mRNA abundance), fundamentally challenging a simplistic, universal m6A effect. Specifically, the observed mRNA decay for highly modified transcripts with high m6A ratios aligns with the known function of m6A readers like YTHDF2, which facilitate recruitment of the CCR4-NOT complex for deadenylation and degradation^49^. In contrast, the increased abundance of transcripts with lower m6A stoichiometry could be mediated by other m6A binding proteins, such as YTHDC1^51^ or IGF2BPs^52^, implicated in enhancing mRNA stability^53^. This bimodal rheostat model provides a refined paradigm for epitranscriptomic regulation, suggesting m6A acts as a dynamic, tunable modulator of gene expression rather than a simple binary switch. While the precise interplay of competing m6A reader proteins and their cofactors, influenced by RNA sequence and stoichiometry, remains an active area of investigation^53,54^, our study provides the first comprehensive analysis in the heart to link specific m6A mRNA motifs at a single-nucleotide level with these divergent aspects of gene processing. This principle is likely to extend beyond cardiogenesis, influencing gene dosage in other developmental and physiological processes where precise control of mRNA abundance is paramount.

We further unveil a novel post-transcriptional m6A-YY1 regulatory axis that is indispensable for orchestrating the critical cardiac metabolic switch. YY1 is a well-established transcription factor known for its essential roles in cardiac progenitor cell lineage commitment and maintenance^55–57^, with its deletion leading to severe cardiac defects^58^. However, YY1 exerts pleiotropic effects on cellular metabolism, functioning both as an activator and repressor of metabolic pathways like glycolysis, the pentose phosphate pathway, OXPHOS, and mitochondrial function, depending on the cellular context^41–43,59–61^. Importantly, its specific regulatory mechanisms during mid-gestation, a period of dynamic metabolic remodeling in the developing heart, have remained largely underexplored. Our integrated multi-omics analyses now demonstrate that Mettl14-m6A epitranscriptomics precisely controls YY1 protein levels via a novel post-transcriptional mechanism, without altering its mRNA abundance. This m6A-mediated regulation of YY1, by mitigating its suppressive effect on ETC genes, consequently, unveils a novel function for YY1 during mid-to-late cardiogenesis. This finding provides a new layer to the complex regulatory network governing transcription factor activity, expanding how epitranscriptomics indirectly controls metabolic reprogramming.

Our study is further distinguished by the development and application of a pioneering CRISPR-based hiPSC platform for targeted epitranscriptomic editing, representing a critical advance in deciphering mRNA modification functions. This cutting-edge tool addresses a long-standing need, as traditional global gain- or loss-of-function approaches lack the spatial and temporal precision to dissect the dynamic, gene-specific regulatory roles of RNA modifications^62–64^. Adapting the CRISPR/Cas13 system^65^, our platform uses dCas13 fusion constructs to introduce m6A marks with remarkable site-specificity at any desired transcript site within hiPSC-derived tissues, providing an unprecedented tool for precise RNA manipulation. This platform was instrumental in directly validating the m6A-YY1 axis in differentiated hiPSC-CMs, demonstrating gene-specific effects as targeted m6A methylation of *ACTB* 3’UTR led to mRNA downregulation, while the same modification in *YY1* coding and 3’UTR resulted in protein downregulation exclusively, underscoring the nuanced nature of m6A-mediated regulation.

Critically, our platform revealed that the regulatory outcome of targeted m6A editing is highly sensitive to both the precise editing site location and the cellular context. Applying identical editing strategies in undifferentiated hiPSCs versus differentiated hiPSC-CMs yielded strikingly different functional outputs. For example, while targeted m6A methylation consistently suppressed YY1 protein in hiPSC-CMs, applying the same editing to undifferentiated hiPSCs did not increase gene and protein expression; instead, it elicited region-specific, differential effects. This powerful observation provides compelling evidence that m6A signals are dynamically interpreted by the cellular environment and developmental stage, constituting an actively interpreted code that yields adaptive and nuanced regulatory outcomes, rather than a fixed, universal response. Our hiPSC-based platform therefore represents an invaluable resource for future mechanistic studies, unlocking new avenues for understanding RNA-based regulation in human health and disease by addressing key research questions concerning the mRNA-protein disconnect, the role of stoichiometry, and cell-type specificity of m6A-mediated regulation in isogenic hiPSC-derived tissues.

In conclusion, our study fundamentally redefines the role of m6A epitranscriptomics as a sophisticated, context-dependent regulatory layer. By uncovering a stoichiometry-governed bimodal code and a novel post-transcriptional m6A-YY1 axis, enabled by unprecedented technological approaches, we reveal how RNA modifications precisely orchestrate complex cellular programs. These findings establish broadly applicable molecular principles for gene regulation in development, stem cell biology, and disease etiology, and provide a conceptual and technical framework for future targeted epitranscriptomic interventions across diverse tissues.

## Limitations of the study

Despite our comprehensive analysis, certain limitations warrant consideration. Our study primarily focused on m6A-dependent effects of Mettl14. While the profound global suppression of m6A modifications in M14-KO hearts, as revealed by nanopore m6A-Seq, strongly implicates m6A-mediated regulation, we cannot entirely exclude potential m6A-independent functions of Mettl14. Furthermore, other regulatory layers, such as non-coding RNAs or additional transcription factors, may also contribute to the observed OXPHOS gene dysregulation. Another challenge involved accurately quantifying translational efficiency; while our proteomics data correlated protein abundance with m6A motifs, the level of steady-state protein abundance captured by mass spectrometry is often obscured by protein turnover, degradation, and post-translational regulation. While ribosome profiling is a potential approach, it was unfeasible due to the minimal tissue yield from E12.5 mouse hearts, and the substantially larger sample inputs required. Future advancements in single-cell and low-input protein synthesis methodologies will be critical to fully unravel these nuanced regulatory effects.

## Methods

### Experimental Animals

All animals were housed at the Johns Hopkins Medical Institute institutional animal facility, and experimental procedures were performed in accordance with experimental protocol approved by local Institutional Animal Care and Use Committees (IACUC). *Mettl14 flox/flox* mice were kindly provided by Dr. Chuan He (University of Chicago)^7^ and were bred with non hypomorphic *Nkx2.5-Cre* transgenic mice (Harvey Lab)^32^ to generate *Nkx2.5-cre; Mettl14 flox/flox* (mutant) and littermate *Cre* negative controls. Embryonic whole heart /ventricles were collected at embryonic day E12.5 and E15.5 as indicated. Dissections were performed in ice-cold PBS under a stereoscopic microscope as previously^66^. Ventricles were either snap-frozen in liquid nitrogen for RNA or protein isolation, fixed for histological analyses, or processed immediately for single-cell dissociation. Both sexes were used in all experimental assays.

### Cell lines

All cells were maintained at 37 °C and 5% CO₂. Human induced pluripotent stem cells (hiPSCs) (WTC11 line, Conklin lab, UCSF) were cultured on Geltrex (Gibco, A1413302)-coated plates in Essential 8 medium (Gibco, A1517001) and passaged between P50–P60 for experiments. CM differentiation was performed by temporal modulation of Wnt signaling as previously described^66^. Briefly, at ∼90% confluency (Day 0), hiPSCs were treated with CHIR99021 (Tocris, 4423) in RPMI media (Gibco, 11875119) supplemented with B27 minus insulin (Gibco, A1895601) for 48 hours, followed by RPMI B27 minus insulin only for 24 hours. From Day 3 to Day 5, cells were treated with *endo* IWR-1 (Tocris, 3532) in RPMI B27 minus insulin. Media were then changed to RPMI B27 minus insulin for 48 h, and RPMI supplemented with B27 (Gibco, 17504044) from Day 7 onward. Beating cardiomyocytes typically emerged between Day 7–9 and were used for downstream assays, immunostaining, RNA/protein extraction, or functional studies.

### m6A LC-MS/MS analysis

For the detection and quantification of m^6^A in mRNA, 500 ng of poly(A) mRNA was denatured at 70°C for 5 minutes followed by digestion to nucleotides using 20 units of S1 Nuclease (Thermo Scientific) in S1 Nuclease buffer for 2 hours at 37°C in 25 μL reactions. Nucleotides were then dephosphorylated to nucleosides by the addition of 2 units of Fast Alkaline Phosphatase (NEB) in FastAP reaction buffer for 1 hour at 37°C. 5 μL of the filtered solution was analyzed by LC-MS/MS. The separation of nucleosides was performed using an Agilent 1290 UHPLC system with a C18 reversed-phase column (2.1 × 50 mm, 1.8 m). The mobile phase A was water with 0.1% (v/v) formic acid and mobile phase B was methanol with 0.1% (v/v) formic acid. Online mass spectrometry detection was performed using an Agilent 6470 triple quadrupole mass spectrometer in positive electrospray ionization mode. Quantification of each nucleoside was accomplished in dynamic multiple reaction monitoring (dMRM) mode by monitoring the transitions of 268→136 (A), 282→150 (m^6^A). The amounts of A and m^6^A in the samples were quantified using corresponding calibration curves generated with pure standards.

### In utero echocardiography

Pregnant *Mettl14 flox/flox* female carrying E15.5 embryos were anesthetized and in utero echocardiography was performed as previously described^67^. Embryo isolation and genotyping was performed as above.

### Western Blotting

Embryonic ventricles or hiPSC-derived CMs were lysed in 1x Cell Lysis Buffer (Cell Signaling Technology, 9803) supplemented with 1mM PMSF. Lysates were briefly sonicated on ice, followed by centrifugation at 14,000 g for 10 min at 4 °C. Protein concentration was measured using the Pierce BCA Protein Assay Kit (Thermo Fisher Scientific, 23227). Equal amounts of protein (10-15 µg per sample were mixed with 2x SDS loading buffer (Invitrogen, LC2676), boiled for 5 min, and resolved on 4-20% TGX precast gels (Bio-Rad, 4561094). Proteins were transferred to 0.2 µm Nitrocellulose membranes (Bio-Rad, 1704271) using Trans-Blot Turbo Transfer System. Following transfer, membranes were stained with Revert Total Protein Stain (LI-COR, 926-11010) for normalization, destained according to manufacturer’s instructions, and then blocked by Intercept (TBS) Blocking Buffer (LI-COR, 927-60001) for 1 hour at room temperature (RT). Membranes were incubated overnight at 4 °C with primary antibodies diluted in blocking buffer, including METTL14 (Proteintech, 26158-1-AP, 1:2000) and YY1 (Cell Signaling Technologies, 63227S, 1:1000). After washing, membranes were incubated with IRDye 800CW secondary antibodies (LI-COR, 925-32211, 1:5,000). Fluorescent signals were imaged using a LI-COR Odyssey system, and protein abundance was quantified using Image Studio, with total protein normalization applied.

### Quantitative Real-Time PCR (qPCR)

Total RNA was isolation from embryonic ventricles or hiPSC-derived CMs using either TRIzol or RNeasy kits (Qiagen, Mini 74104 and Micro 74004) according to manufacturers’ protocols. RNA concentration and purity were determined by NanoDrop spectrophotometry. cDNA was synthesized from 0.5-1 µg RNA using High-Capacity cDNA Reverse Transcription Kit (Applied Biosystems, 4374966). qPCR was performed using SYBR Select Master Mix (Applied Biosystems, 4472908) on a QuantStudio 5 Real-Time PCR System (Applied Biosystems, 384-well format). Reactions were run in technical duplicates.

Gene expression levels were calculated using the ΔΔCt method and normalized to beta 2 microglobulin (B2M). Data are presented as fold change relative to control samples. Primer sequences used for amplification were purchased from Integrated DNA Technologies, sequences listed in oligonucleotide table.

### mtDNA content analysis

Genomic DNA was isolated using Quick-DNA Microprep Kit (Zymo Research, D3020). qPCR was performed using SYBR Select Master Mix (Applied Biosystems, 4472908), adding 10ng of template DNA per reaction and using mtDNA target mt-Nd1 (mouse) and mt-ND1 (human) and nuclear-encoded reference gene B2m (mouse) or Rpl32 (human).

### Global m6A quantification

Global m6A quantification: Global m6A levels were measured using the Epiquik m6A RNA Methylation Quantification Kit (Colorimetric) (EPIGENTEK, P-9005-48), according to manufacturer’s instructions. Briefly, 50ng of total RNA were isolated from in vitro CMs was used as starting material. RNA was bound to assay wells, incubated with capture and detection antibodies, and colorimetric signals were quantified at OD450 nm using a microplate reader.

### Histology, immunostaining, EdU, and TUNEL

Animals were euthanized by isoflurane anesthesia followed by cervical dislocation and removal of whole hearts (for sectioning) or ventricles only (all other purposes) at embryonic day E12.5 and E15.5, as indicated. Embryonic hearts/ventricles were fixed in 4% paraformaldehyde (PFA), overnight, cryoprotected in 30% sucrose, embedded in OCT, and sectioned at 6 µm.

For histological analyses, sections were submitted to the Oncology Tissue & Imaging Services (OTIS) Core lab at Johns Hopkins Medicine for hematoxylin and eosin staining to assess morphology.

For tissue immunofluorescence, sections were washed thrice in PBS for rehydration, permeabilized in 0.2% Triton X-100 in PBS (PBS-T) thrice 8 minutes each, then blocked for 1hour at RT in 1% BSA. Primary antibodies were applied overnight at 4 °C, including cardiac troponin T (cTnT) (Abcam, ab8295, 1:500 in 1% BSA) and phospho-Histone H3 (Millipore, 06-570, 1:500 in 1% BSA). After PBS washes, sections were incubated with Alexa Fluor 488/594/647-conjugated secondary antibodies (Abcam, 1:500 in 1% BSA) for 1hour at RT, washed thrice, Hoechst 33342 (Thermo Fisher, 62249; 1:2,000 in PBS) for nuclei staining, followed by mounting with ProLong Diamond Antifade (Thermo Fisher, P36962).

For immunostaining of cultured cells, hiPSC-derived CMs were seeded on plastic coverslips in 24-well plates. Cells were fixed in 4% PFA for 15 min at RT with gentle shaking, washed in PBS, and permeabilized in 0.2% PBS-T. Blocking was performed in 1% BSA in PBS for 1hour, followed by overnight incubation with primary antibodies at 4 °C. After 3 PBS washes, cells were incubated with secondaries for 1hour at RT, washed again, counterstained with Hoechst, washed with PBS, and mounted with coverslips.

EdU proliferation assay: In vitro, cells were incubated with EdU labeling solution (Click-iT EdU 647, Thermo Fisher) for 1 hour at 37 °C before fixation. In vivo, pregnant females received maternal intraperitoneal injection of EdU without disturbing embryos. Embryos were harvested 45 min post-injection, and EdU was detected in cryosections according to manufacturer’s protocol. If immunostaining and EdU detection were combined, secondary antibodies were diluted directly in the EdU reaction cocktail the following day.

TUNEL apoptosis assay: Click-iT Plus TUNEL assay (Thermo Fisher) was performed on heart cryosections prior to blocking, followed by immunostaining if required.

Imaging was performed using a Leica SP8 confocal microscope, and analysis was conducted in Fiji by blinded investigators.

### Single cell RNA sequencing

Hearts at E12.5 were dissected, atria removed, and tail tissue isolated for genotyping. Ventricles were carefully dissociated in TryPLE express (Gibco, 12604013) at 37 °C for 15 min. Following digestion, cells were neutralized with Dulbecco’s modified Eagle’s medium (DMEM) (Corning, 10-017-CV) supplemented with 5% fetal bovine serum (FBS), centrifuged, and resuspended in cold PBS containing 5% FBS. Cell suspensions were passed through a 70 µm cell strainer, centrifuged again, and resuspended in PBS+5% FBS on ice.

Cell viability ≥ 98% was confirmed before downstream sequencing processing. Cell encapsulation, reverse transcription, library preparation, sequencing, and Cell Ranger (10X Genomics) were performed by the institutional single cell & transcriptomics core facility using 10X Genomics Chromium Single Cell 3’ HT platform. Libraries were sequenced on an Illumina NovaSeq S4 flow cell (200-cycle). FASTQ files were processed using Cell Ranger and aligned to the Mus musculus mm39 reference genome.

Filtered feature-barcode matrices were combined and analyzed using the Seurat (v5.2). The combined Seurat object was pre-processed through QC (percent mitochondrial reads < 25), normalization, feature selection, and scaling. 8836 controls and 8367 M14-KOs were captured and subjected to PCA, clustering, and UMAP visualization. Cardiomyocytes clusters were identified by expression of sarcomeric markers (Ttn, Myh6, Myl2). Differential gene expression between control and M14-KO cardiomyocytes were computed using FindMarkers (Wilcoxon rank-sum test), with genes considered significant at adjusted p ≤ 0.05 (Benjamini-Hochberg) and expression in ≥ 10% of cells, |Log_2_FC| ≥ 0.25.

### Nanopore Direct RNA sequencing

Total RNA was isolated from embryonic hearts at E12.5 (atria excised) using Micro RNeasy kit (Qiagen). RNA integrity was confirmed prior to library preparation. Direct RNA libraries were generated using Oxford Nanopore Direct RNA Sequencing Kit and sequenced on a PromethION platform. Base calling was performed using Dorado (Oxford Nanopore Technologies) with the super accuracy model, in parallel with modification-aware models for detecting modified adenosines, and reads were aligned to the Mus musculus mm39 reference genome using minimap2 with splice-aware parameters. Modkit (Oxford Nanopore Technologies) was used to generate per-site modification summary statistics. Sites with modification probability ≥ 0.9 were considered m6A-modified, whereas positions below this threshold were treated as canonical A. Only sites supported by ≥10 reads in at least three biological replicates per condition were retained. Site-level percent modification was calculated per sample, and methylation difference were reported as delta percent modification = Control – M14-KO.

### Proteomics (Label-Free Quantitative Mass Spectrometry)

Ventricles collected from E12.5 hearts were processed for proteomics using the iST 8x Sample Preparation Kit (PreOmics, P.O.00001). Briefly, embryonic ventricles were reduced and alkylated in iST lysis buffer at 95 °C for 10 min, followed by sonication. Then the lysates were subjected to on-column digestion with trypsin/Lys-C at 37 °C. Peptides were acidified, transferred to cartridge, washed, eluted to axygen 96 well plate (Corning, PMI110-07A), vacuum-dried, sealed, and shipped out overnight for LC-MS/MS.

Peptides were loaded onto a Dionex RSLC Ultimate 300 (Thermo Scientific) coupled to an Orbitrap Exploris 480 Mass spectrometer (Thermo Fisher Scientific) for separation as described^68^. Raw proteomics files were analyzed using FraqPipe to obtain protein identifications and label-free quantification (LFQ) values with in-house parameters (variance-stabilizing normalization and k-nearest neural network for imputation). Prior to statistical analysis, common contaminants, reverse hits, and proteins not consistently identified and quantified across biological replicates within each condition were removed. The LFQ intensities were log-transformed, and missing values were imputed using a Missing Not At Random (MNAR) approach. Differential expression was computed using limma package (R Bioconductor) with empirical Bayes moderation, and proteins with |Log_2_FC|≥0. 25 and adjusted p.mod ≤ 0.05 were considered significant.

### Adenovirus Infection and siRNA Transfection

Adenoviral infection: For loss-of-function experiments, hiPSC-derived CMs were replated on 0.1% gelatin-coated wells and infected at ∼70–80% confluency. Cells were incubated with adenovirus encoding either GFP control (Ad-Mock) or METTL14 shRNA (Ad-M14sh, purchased from VectorBuilder, #VB230420-1197tda) at a multiplicity of infection (MOI) of 10–50, diluted in culture medium. After 24 hours, cells were either collected for downstream RNA and protein processing or swapped with fresh RPMI+B27 medium. Cells were collected 96 hours post-infection for RNA, protein, immunostaining, or functional assays. Infection efficiency was confirmed by reduction in target transcript/protein levels.

siRNA transfection: For transient knockdown in rescue experiments, Adenovirus infected hiPSC-derived CMs were transfected with siRNA targeting YY1 (siYY1), or non-targeting control siRNA (siNC) using Lipofectamine RNAiMAX (Thermo Fisher Scientific, 13778030) according to the manufacturer’s protocol. Briefly, 12 hours post infection, siRNA (2-5 pmol) and RNAiMAX were diluted separately in Opti-MEM, combined for 10 min incubation at RT, and added to cells dropwise. Cells were harvested 12 hours after transfection for downstream qPCR analysis. Knockdown efficiency confirmed at the protein level.

### m6A RNA immunoprecipitation-qPCR

M6A RNA immunoprecipitation (MeRIP)-qPCR: To assess transcript-specific m6A enrichment, MeRIP-qPCR was performed using the EpiQuik CUT&RUN m6A RNA Enrichment Kit (EPIGENTEK, P-9018-24). 15-20 µg total RNA was used for each immunoprecipitation reaction, while 400 ng RNA was reserved as input control. RNA was fragmented, incubated with anti-m6A antibody-conjugated beads, washed, and eluted. RNA from IP and input fractions was reverse-transcribed and subjected to qPCR. Relative enrichment was first calculated as ΔCt = Ct(m6A-IP) - Ct(Input). Fold enrichment in M14-KD samples relative to control was then calculated using: 2^-[(M14-KD ΔCt) - (Control ΔCt)].

Measurement of in vitro oxygen consumption rates (OCR) and extracellular acidification rate (ECAR) hiPSC-derived CMs were plated on Agilent Seahorse XFe96 cell culture microplates (Agilent) at a density of 50,000 cells per well. After 48 hours, cells were infected with either Ad-Mock or Ad-M14sh to induce m6A loss *in vitro*. Mitochondrial respiration was assessed 96 hours post-infection using Agilent Seahorse XFe96 Analyzer and Agilent Seahorse Mito Stress Test Kit (Agilent) following manufacturer’s instructions. Oligomycin (Sigma–Aldrich, 75351-5MG) was loaded into port A at a final concentration of 1 µM. Carbonyl cyanide 4-(trifluoromethoxy) phenylhydrazone (FCCP; Sigma–Aldrich, C2920-10MG) was loaded into port B at a final concentration of 2 µM. Rotenone (Sigma–Aldrich, R8875-1G) and antimycin A (Sigma–Aldrich, A8674-25MG) were loaded into port Cat final concentrations of 0.5 µM. The assay consisted of 12 measurement cycles, including 3 baseline cycles, followed by 3 cycles after each compound injection. OCRs and ECAR were analyzed using Wave software (Agilent) and normalized by the number of nuclei by Hoechst staining. ≥5 technical replicates and ≥3 biological replicates were included.

### Substrate Utilization Assay

Individual complex-linked respiration was evaluated using substrate utilization assay, as described previously^69^. At the day of experiment, cells will be swapped into media containing complex-specific substrates (CI-linked: 10 mM pyruvate, 2 mM malate; CII-linked: 5 mM succinate, 10 µM rotenone; CIII-mediated: 10 mM glycerol-3-phosphate/rotenone(G3P), 2 mM malate, 2 µM rotenone; CIV-linked: 10 mM ascorbate, 100 µM TMPD, 2 µM antimycin A), followed by treatment with digitonin (port A; final concentration: 25 µg/mL) to selectively permeabilize the plasma membrane while preserving mitochondrial integrity, for basal respiration measurements (State 4). ADP (port B; final concentration: 4mM) was then injected to stimulate State 3, ADP-stimulated respiration. The assay consisted of 9 measurement cycles, including 3 baseline cycles, followed by 3 cycles after digitonin and ADP injection, respectively. State 3 and state 4 OCR values normalized by CyQUANT Cell Proliferation Assays (Invitrogen, C7026). Respiratory Control Ratio (RCR) was calculated as the ratio of state 3 to state 4 respiration. ≥5 technical replicates and ≥3 biological replicates were included for each complex.

### Gene ontology and Transcription Factor (TF) analysis

Gene Ontology (GO) enrichment analysis of differentially expressed genes or proteins from the indicated comparisons was performed using Enrichr^70^ against the GO biological Process 2021 database. Adjusted P-values were calculated using the Benjamini–Hochberg method, and only top nonrepetitive significant pathways (adjusted P < 0.05) were reported. Gene ratio was defined as the number of genes in the input list divided by the total number of genes assigned to a given GO term. TF enrichment analysis was performed using the ENCODE and ChEA Consensus TFs from ChIP-X database implemented in Enrichr, by inputting differentially expressed genes. Enriched transcription factors (e.g., BRCA1, TAF1, YY1, PML, NFYB) were ranked according to adjusted P value and combined enrichment score.

### CRISPR-based targeted m6A epitranscriptomic editing

For enhancer targeted epitranscriptomics editing experiments, a hiPSC line was generated from the parent line (WTC11). Specifically, the deactivated Cas13b fused with Mettl3-NLS (cloned from pCMV-dCas13nls plasmid, Addgene #155366) under the constitutive expression of a CAG promoter, was knocked in the AAVS locus using CRISPR/Cas9^66^. sgRNAs were cloned in the plasmid carrying crRNA scaffold compatible with Cas13b (pC0043-PspCas13b crRNA backbone, Addgene #103854). Two different gRNA plasmids targeting the A1296 and A1513 regions of *YY1* and one gRNA targeting the A1216 *ACTB* 3’UTR region were used. The empty vector was used as control. Lipofectamine Stem reagent (ThermoFisher) was used to transfect day 7 hiPSC-CMs according to the manufacturer’s instructions. Cells were isolated four days post-transfection.

### Quantification and statistical analysis

Statistical analyses were performed using GraphPad Prism 10 (GraphPad Software Inc). All studies in cultured cells were performed using at least three sets of independent experiments. For in vivo studies, at least five animals in each group were analyzed. For all experiments, the numbers of samples used are specified in the respective figure legends. The Shapiro–Wilk test was performed to assess normal distribution, and parametric or non-parametric tests were used as appropriate for two or more group analyses as reported in the respective figure legends.

## Methods – Tables

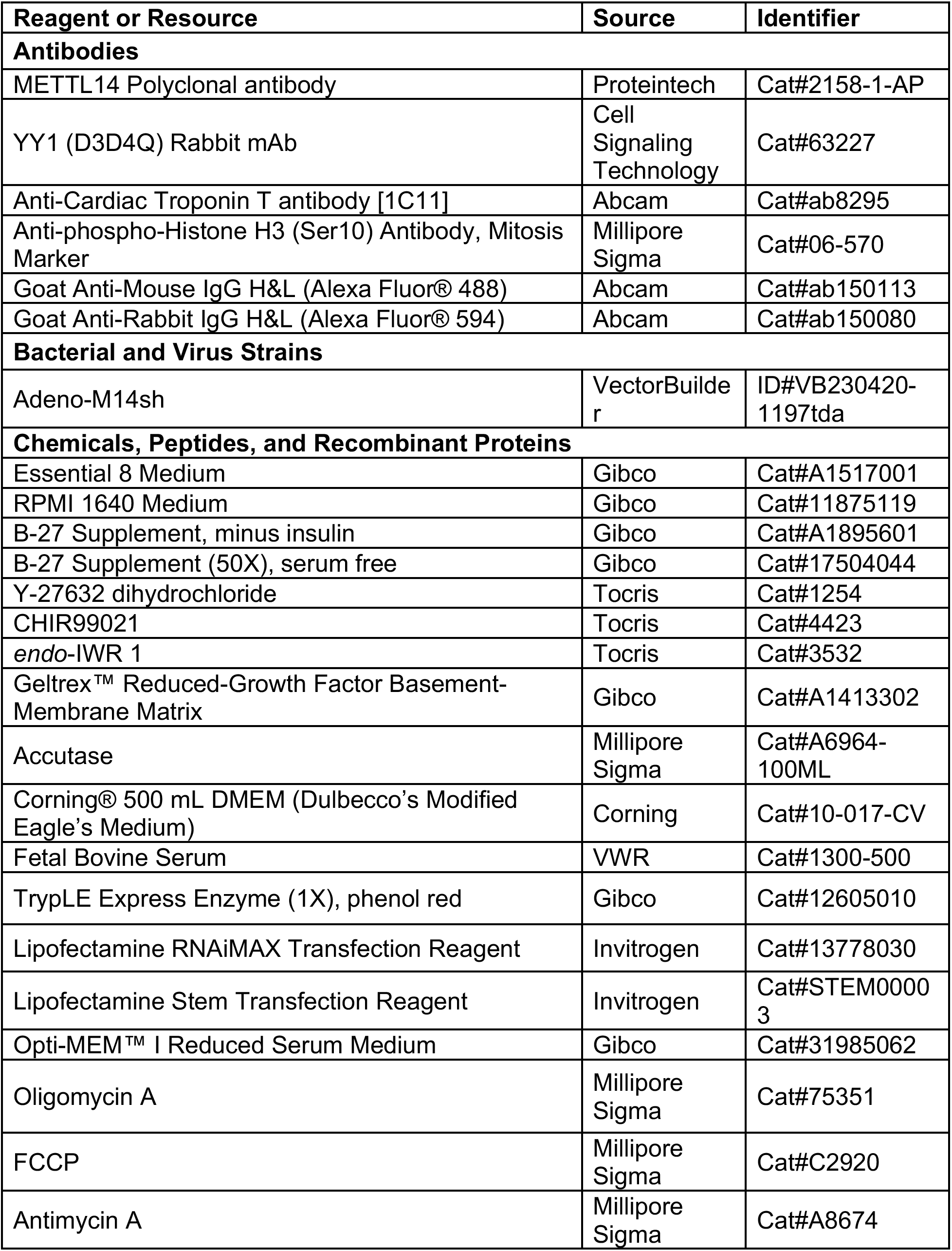
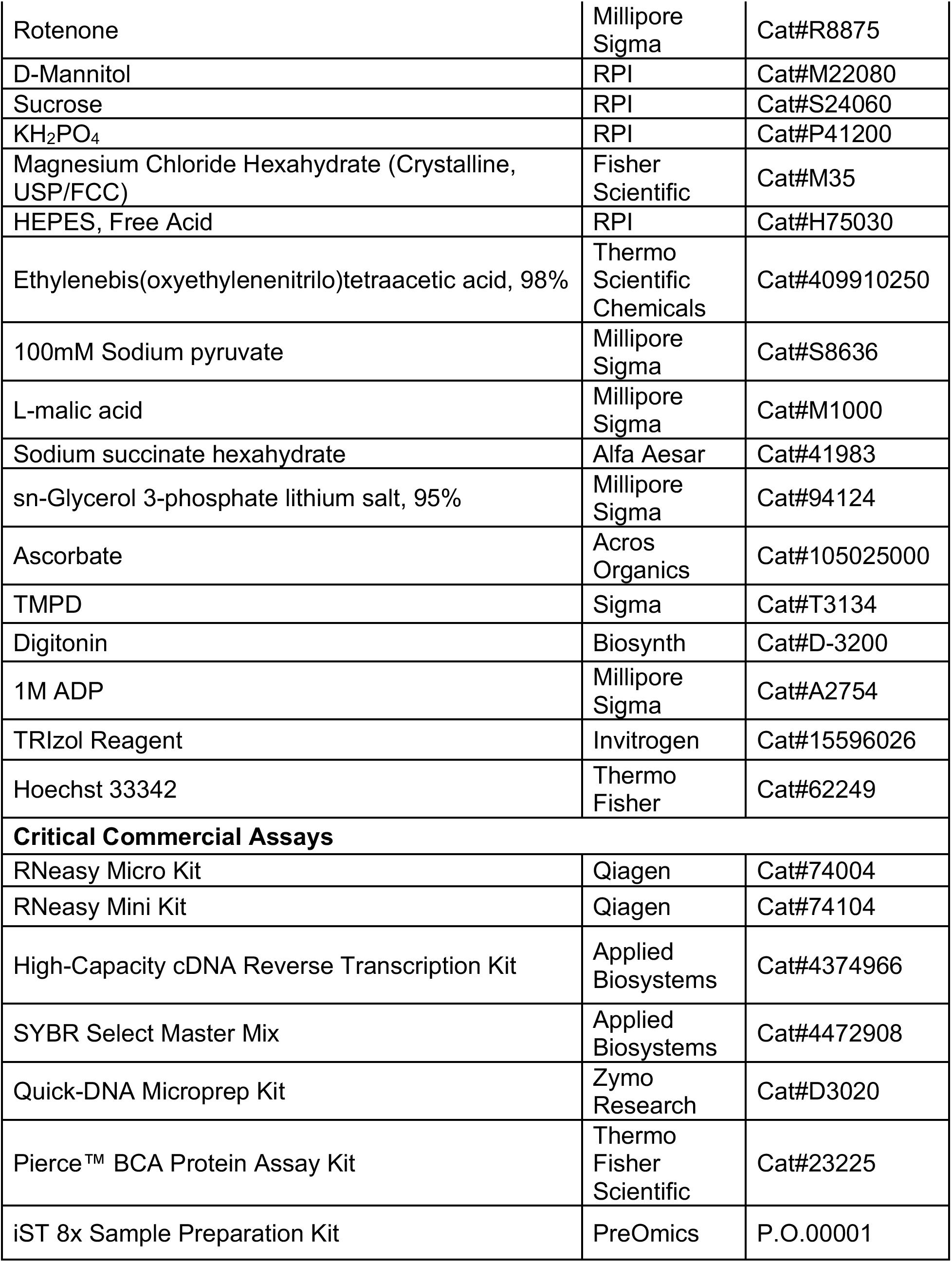
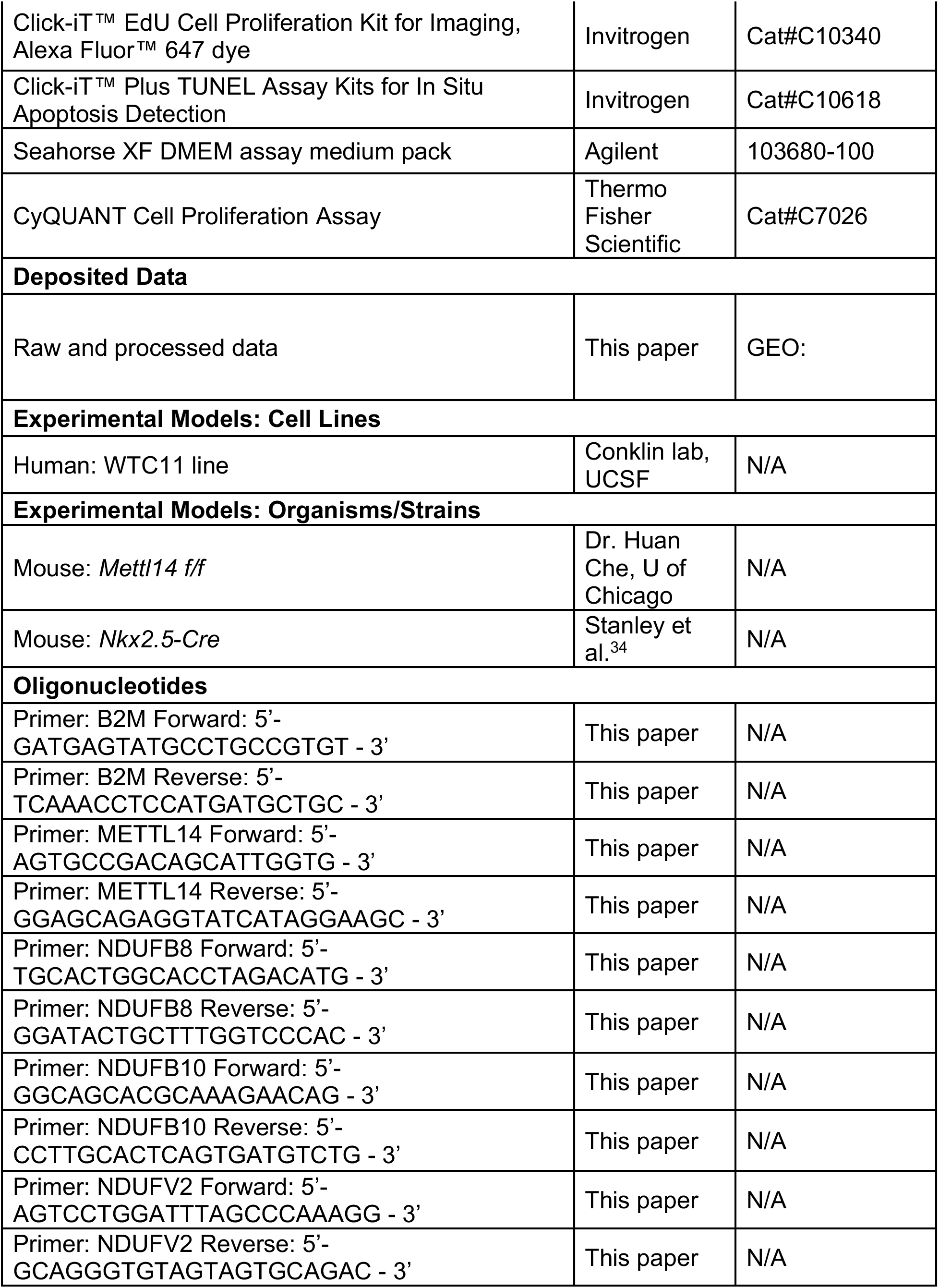
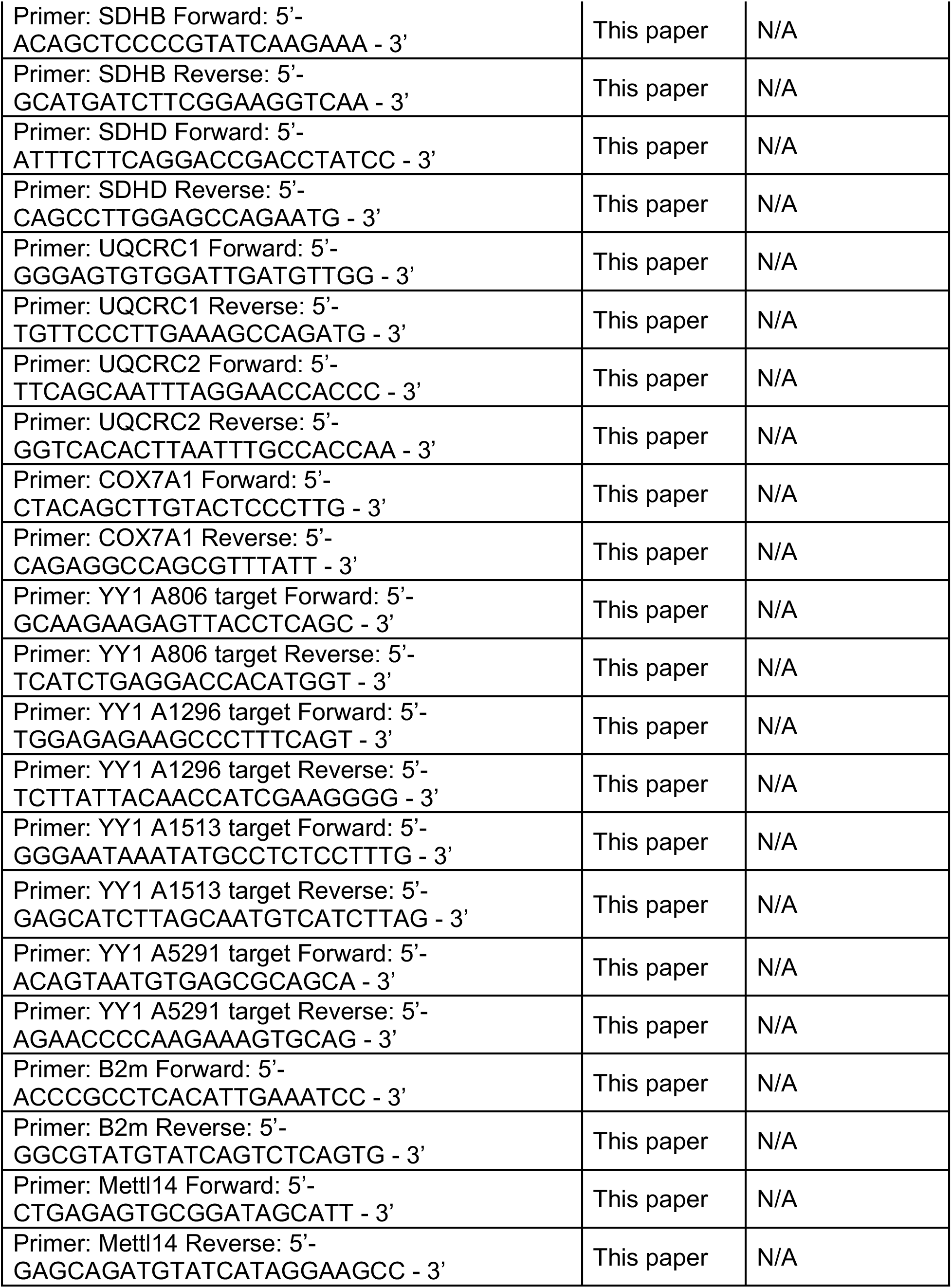
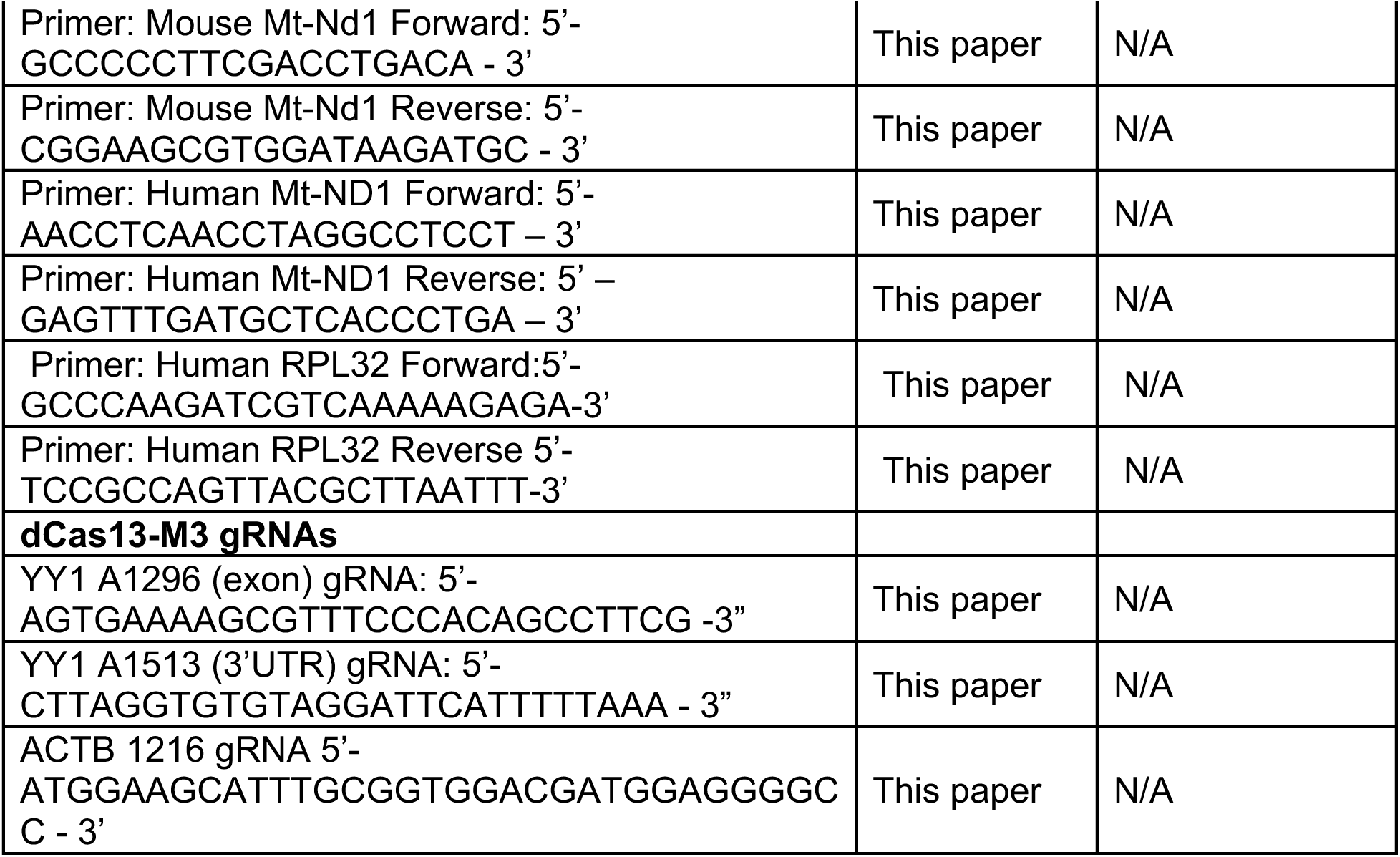

| **Recombinant DNA** |  |  |
| --- | --- | --- |
| siYY1 | IDT | Design ID#hs.Ri.YY1.13.2 |
| Negative Control DsiRNA | IDT | Cat#51-01-14-03 |
| **Software and algorithms** |  |  |
| Seurat v5 | Hao et al.<sup>71</sup> | https://satijalab.org/seurat |
| Dorado | Oxford Nanopore Technologies | https://github.com/nanoporetech/dorado |
| Modkit | Oxford Nanopore Technologies | https://github.com/nanoporetech/modkit |
| FragPipe | Yu et al.<sup>72</sup> | https://github.com/Nesvilab/FragPipe |
| Limma | Bioconductor | https://bioconductor.org/packages/release/bioc/html/limma.html |

## Acknowledgements

The authors would like to thank all members of the Tampakakis lab for the insightful comments and recommendations. We would also like to thank Dr. Chuan He (University of Chicago) for kindly providing the Mettl14 fl/fl mice and Dr. Navid Koleni for assisting with the generation of adenoviral vectors.

E.T. is supported by grants from NHLBI (HL-145135), The Johns Hopkins University Catalyst Award, and MSCRF awards (2023-MSCRFL-5984 and 2024-MSRFD-6393. This work was also supported in part by NIH grants (R01 HG010538 to WT, R01HL165729 to SMC; K99HL168075 to NS).

## Author contributions

S.L. designed, carried out, supervised this work and wrote the manuscript. HG, SB and NS assisted with experimental work. EY and SS assisted with the proteomics analyses. KB assisted with m6A mass-spectrometry. CH, QG and WT assisted with all nanopore-Sequencing experiments and associated bioinformatics analyses. SMC assisted with all ETC complex analyses. ET designed, supervised this work, assisted with experimental work, and wrote the manuscript. All authors have read and approved the final version of the manuscript.

## Competing interests

Dr. Winston Timp has two patents (8,748,091 and 8,394,584) licensed to Oxford Nanopore Technologies (ONT) none of which are related to this work. He also received reimbursement for travel, accommodation, and/or conference fees to speak at events organized by ONT. All other authors have no competing interests to declare.

## Notes

### Competing Interest Statement

The authors have declared no competing interest.

## References

1 Wang, G., Wang, B. & Yang, P. Epigenetics in Congenital Heart Disease. J Am Heart Assoc 11, e025163 (2022). 10.1161/JAHA.121.025163

2 Miyamoto, M., Gangrade, H. & Tampakakis, E. Understanding Heart Field Progenitor Cells for Modeling Congenital Heart Diseases. Curr Cardiol Rep 23, 38 (2021). 10.1007/s11886-021-01468-5

3 Rowton, M., Guzzetta, A., Rydeen, A. B. & Moskowitz, I. P. Control of cardiomyocyte differentiation timing by intercellular signaling pathways. Semin Cell Dev Biol (2021). 10.1016/j.semcdb.2021.06.002

4 Christoffels, V. & Jensen, B. Cardiac Morphogenesis: Specification of the Four-Chambered Heart. Cold Spring Harb Perspect Biol 12 (2020). 10.1101/cshperspect.a037143

5 Geula, S. et al. Stem cells. m6A mRNA methylation facilitates resolution of naive pluripotency toward differentiation. Science 347, 1002–1006 (2015). 10.1126/science.1261417

6 Zhang, C. et al. m(6)A modulates haematopoietic stem and progenitor cell specification. Nature 549, 273–276 (2017). 10.1038/nature23883

7 Weng, H. et al. METTL14 Inhibits Hematopoietic Stem/Progenitor Differentiation and Promotes Leukemogenesis via mRNA m(6)A Modification. Cell stem cell 22, 191–205 e199 (2018). 10.1016/j.stem.2017.11.016

8 Sendinc, E. & Shi, Y. RNA m6A methylation across the transcriptome. Mol Cell 83, 428–441 (2023). 10.1016/j.molcel.2023.01.006

9 Wang, S. et al. Dynamic regulation and functions of mRNA m6A modification. Cancer Cell Int 22, 48 (2022). 10.1186/s12935-022-02452-x

10 Shi, H., Wei, J. & He, C. Where, When, and How: Context-Dependent Functions of RNA Methylation Writers, Readers, and Erasers. Mol Cell 74, 640–650 (2019). 10.1016/j.molcel.2019.04.025

11 Gatsiou, A. & Stellos, K. RNA modifications in cardiovascular health and disease. Nat Rev Cardiol 20, 325–346 (2023). 10.1038/s41569-022-00804-8

12 Huang, H., Weng, H. & Chen, J. The Biogenesis and Precise Control of RNA m(6)A Methylation. Trends Genet 36, 44–52 (2020). 10.1016/j.tig.2019.10.011

13 Workman, R. E. et al. Nanopore native RNA sequencing of a human poly(A) transcriptome. Nature methods 16, 1297–1305 (2019). 10.1038/s41592-019-0617-2

14 Garalde, D. R. et al. Highly parallel direct RNA sequencing on an array of nanopores. Nature methods 15, 201–206 (2018). 10.1038/nmeth.4577

15 Diensthuber, G. & Novoa, E. M. Charting the epitranscriptomic landscape across RNA biotypes using native RNA nanopore sequencing. Mol Cell 85, 276–289 (2025). 10.1016/j.molcel.2024.12.014

16 Schwartz, S. et al. Perturbation of m6A writers reveals two distinct classes of mRNA methylation at internal and 5’ sites. Cell reports 8, 284–296 (2014). 10.1016/j.celrep.2014.05.048

17 Edwards, W. et al. Quantitative proteomic profiling identifies global protein network dynamics in murine embryonic heart development. Developmental cell 58, 1087–1105 e1084 (2023). 10.1016/j.devcel.2023.04.011

18 Gu, Y. et al. Multi-omics profiling visualizes dynamics of cardiac development and functions. Cell reports 41, 111891 (2022). 10.1016/j.celrep.2022.111891

19 An, Y. & Duan, H. The role of m6A RNA methylation in cancer metabolism. Mol Cancer 21, 14 (2022). 10.1186/s12943-022-01500-4

20 Li, Q. et al. HIF-1alpha-induced expression of m6A reader YTHDF1 drives hypoxia-induced autophagy and malignancy of hepatocellular carcinoma by promoting ATG2A and ATG14 translation. Signal Transduct Target Ther 6, 76 (2021). 10.1038/s41392-020-00453-8

21 Batista, P. J. et al. m(6)A RNA modification controls cell fate transition in mammalian embryonic stem cells. Cell stem cell 15, 707–719 (2014). 10.1016/j.stem.2014.09.019

22 Liu, J. et al. A METTL3-METTL14 complex mediates mammalian nuclear RNA N6-adenosine methylation. Nat Chem Biol 10, 93–95 (2014). 10.1038/nchembio.1432

23 Yoon, K. J. et al. Temporal Control of Mammalian Cortical Neurogenesis by m(6)A Methylation. Cell 171, 877–889 e817 (2017). 10.1016/j.cell.2017.09.003

24 Gu, Y., Song, Y., Pan, Y. & Liu, J. The essential roles of m(6)A modification in osteogenesis and common bone diseases. Genes Dis 11, 335–345 (2024). 10.1016/j.gendis.2023.01.032

25 Yang, X. et al. m(6)A Methylases Regulate Myoblast Proliferation, Apoptosis and Differentiation. Animals (Basel) 12 (2022). 10.3390/ani12060773

26 Xie, S. J. et al. Dynamic m(6)A mRNA Methylation Reveals the Role of METTL3/14-m(6)A-MNK2-ERK Signaling Axis in Skeletal Muscle Differentiation and Regeneration. Front Cell Dev Biol 9, 744171 (2021). 10.3389/fcell.2021.744171

27 Han, Z. et al. ALKBH5-mediated m(6)A mRNA methylation governs human embryonic stem cell cardiac commitment. Mol Ther Nucleic Acids 26, 22–33 (2021). 10.1016/j.omtn.2021.05.019

28 Wang, S. et al. Differential roles of YTHDF1 and YTHDF3 in embryonic stem cell-derived cardiomyocyte differentiation. RNA Biol 18, 1354–1363 (2021). 10.1080/15476286.2020.1850628

29 Kannan, S. et al. Trajectory reconstruction identifies dysregulation of perinatal maturation programs in pluripotent stem cell-derived cardiomyocytes. Cell reports 42, 112330 (2023). 10.1016/j.celrep.2023.112330

30 Galdos, F. X. et al. devCellPy is a machine learning-enabled pipeline for automated annotation of complex multilayered single-cell transcriptomic data. Nature communications 13, 5271 (2022). 10.1038/s41467-022-33045-x

31 Kim, Y. et al. Nanopore direct RNA sequencing of human transcriptomes reveals the complexity of mRNA modifications and crosstalk between regulatory features. Cell Genom 5, 100872 (2025). 10.1016/j.xgen.2025.100872

32 Stanley, E. G. et al. Efficient Cre-mediated deletion in cardiac progenitor cells conferred by a 3’UTR-ires-Cre allele of the homeobox gene Nkx2-5. Int J Dev Biol 46, 431–439 (2002).

33 Miyamoto, M. et al. Cardiac progenitors instruct second heart field fate through Wnts. Proc Natl Acad Sci U S A 120, e2217687120 (2023). 10.1073/pnas.2217687120

34 Zhao, Q., Sun, Q., Zhou, L., Liu, K. & Jiao, K. Complex Regulation of Mitochondrial Function During Cardiac Development. J Am Heart Assoc 8, e012731 (2019). 10.1161/JAHA.119.012731

35 Dorn, G. W., 2nd, Vega, R. B. & Kelly, D. P. Mitochondrial biogenesis and dynamics in the developing and diseased heart. Genes Dev 29, 1981–1991 (2015). 10.1101/gad.269894.115

36 Gibb, A. A. & Hill, B. G. Metabolic Coordination of Physiological and Pathological Cardiac Remodeling. Circulation research 123, 107–128 (2018). 10.1161/CIRCRESAHA.118.312017

37 Baker CN, E. S. Development of aerobic metabolism in utero: requirement for mitochondria function during embryonic and foetal periods. OA Biotechnol. 2, 16–22 (2013).

38 Beutner, G. et al. Coordinated metabolic responses to cyclophilin D deletion in the developing heart. iScience 27, 109157 (2024). 10.1016/j.isci.2024.109157

39 Funk, C. C. et al. Atlas of Transcription Factor Binding Sites from ENCODE DNase Hypersensitivity Data across 27 Tissue Types. Cell reports 32, 108029 (2020). 10.1016/j.celrep.2020.108029

40 Lachmann, A. et al. ChEA: transcription factor regulation inferred from integrating genome-wide ChIP-X experiments. Bioinformatics 26, 2438–2444 (2010). 10.1093/bioinformatics/btq466

41 Chen, F. et al. YY1 regulates skeletal muscle regeneration through controlling metabolic reprogramming of satellite cells. The EMBO journal 38 (2019). 10.15252/embj.201899727

42 Wu, S. et al. Transcription Factor YY1 Promotes Cell Proliferation by Directly Activating the Pentose Phosphate Pathway. Cancer Res 78, 4549–4562 (2018). 10.1158/0008-5472.CAN-17-4047

43 Li, Y. et al. Yin Yang 1 facilitates hepatocellular carcinoma cell lipid metabolism and tumor progression by inhibiting PGC-1beta-induced fatty acid oxidation. Theranostics 9, 7599–7615 (2019). 10.7150/thno.34931

44 Chen, F., Sun, H., Zhao, Y. & Wang, H. YY1 in Cell Differentiation and Tissue Development. Crit Rev Oncog 22, 131–141 (2017). 10.1615/CritRevOncog.2017021311

45 Nan, C. & Huang, X. Transcription factor Yin Yang 1 represses fetal troponin I gene expression in neonatal myocardial cells. Biochemical and biophysical research communications 378, 62–67 (2009). 10.1016/j.bbrc.2008.10.174

46 Sucharov, C. C., Mariner, P., Long, C., Bristow, M. & Leinwand, L. Yin Yang 1 is increased in human heart failure and represses the activity of the human alpha-myosin heavy chain promoter. J Biol Chem 278, 31233–31239 (2003). 10.1074/jbc.M301917200

47 Fan, R. et al. A combined deep learning framework for mammalian m6A site prediction. Cell Genom 4, 100697 (2024). 10.1016/j.xgen.2024.100697

48 Wilson, C., Chen, P. J., Miao, Z. & Liu, D. R. Programmable m(6)A modification of cellular RNAs with a Cas13-directed methyltransferase. Nature biotechnology 38, 1431–1440 (2020). 10.1038/s41587-020-0572-6

49 Du, H. et al. YTHDF2 destabilizes m(6)A-containing RNA through direct recruitment of the CCR4-NOT deadenylase complex. Nature communications 7, 12626 (2016). 10.1038/ncomms12626

50 Wang, X. et al. N(6)-methyladenosine Modulates Messenger RNA Translation Efficiency. Cell 161, 1388–1399 (2015). 10.1016/j.cell.2015.05.014

51 Cheng, Y. et al. N(6)-Methyladenosine on mRNA facilitates a phase-separated nuclear body that suppresses myeloid leukemic differentiation. Cancer Cell 39, 958–972 e958 (2021). 10.1016/j.ccell.2021.04.017

52 Huang, H. et al. Recognition of RNA N(6)-methyladenosine by IGF2BP proteins enhances mRNA stability and translation. Nat Cell Biol 20, 285–295 (2018). 10.1038/s41556-018-0045-z

53 Wei, G. RNA m6A modification, signals for degradation or stabilisation? Biochem Soc Trans 52, 707–717 (2024). 10.1042/BST20230574

54 Murakami, S. & Jaffrey, S. R. Hidden codes in mRNA: Control of gene expression by m(6)A. Mol Cell 82, 2236–2251 (2022). 10.1016/j.molcel.2022.05.029

55 Gregoire, S. et al. Essential and unexpected role of Yin Yang 1 to promote mesodermal cardiac differentiation. Circulation research 112, 900–910 (2013). 10.1161/CIRCRESAHA.113.259259

56 Gregoire, S., Li, G., Sturzu, A. C., Schwartz, R. J. & Wu, S. M. YY1 Expression Is Sufficient for the Maintenance of Cardiac Progenitor Cell State. Stem Cells 35, 1913–1923 (2017). 10.1002/stem.2646

57 Morikawa, Y., Leach, J. & Martin, J. F. Yin-Yang 1, a new player in early heart development. Circulation research 112, 876–877 (2013). 10.1161/CIRCRESAHA.113.300947

58 Beketaev, I. et al. Critical role of YY1 in cardiac morphogenesis. Dev Dyn 244, 669–680 (2015). 10.1002/dvdy.24263

59 Cunningham, J. T. et al. mTOR controls mitochondrial oxidative function through a YY1-PGC-1alpha transcriptional complex. Nature 450, 736–740 (2007). 10.1038/nature06322

60 Blattler, S. M. et al. Defective mitochondrial morphology and bioenergetic function in mice lacking the transcription factor Yin Yang 1 in skeletal muscle. Mol Cell Biol 32, 3333–3346 (2012). 10.1128/MCB.00337-12

61 Varum, S. et al. Yin Yang 1 Orchestrates a Metabolic Program Required for Both Neural Crest Development and Melanoma Formation. Cell stem cell 24, 637–653 e639 (2019). 10.1016/j.stem.2019.03.011

62 Wang, L. et al. METTL14 is required for exercise-induced cardiac hypertrophy and protects against myocardial ischemia-reperfusion injury. Nature communications 13, 6762 (2022). 10.1038/s41467-022-34434-y

63 Dorn, L. E. et al. The N(6)-Methyladenosine mRNA Methylase METTL3 Controls Cardiac Homeostasis and Hypertrophy. Circulation 139, 533–545 (2019). 10.1161/CIRCULATIONAHA.118.036146

64 Jiang, F. Q. et al. Mettl3-mediated m(6)A modification of Fgf16 restricts cardiomyocyte proliferation during heart regeneration. eLife 11 (2022). 10.7554/eLife.77014

65 Rauch, S., He, C. & Dickinson, B. C. Targeted m(6)A Reader Proteins To Study Epitranscriptomic Regulation of Single RNAs. J Am Chem Soc 140, 11974–11981 (2018). 10.1021/jacs.8b05012

66 Htet, M. et al. A transcriptional enhancer regulates cardiac maturation. Nat Cardiovasc Res 3, 666–684 (2024). 10.1038/s44161-024-00484-2

67 Yu, Q., Leatherbury, L., Tian, X. & Lo, C. W. Cardiovascular assessment of fetal mice by in utero echocardiography. Ultrasound Med Biol 34, 741–752 (2008). 10.1016/j.ultrasmedbio.2007.11.001

68 Jani, V. P. et al. Myocardial Proteome in Human Heart Failure With Preserved Ejection Fraction. J Am Heart Assoc 14, e038945 (2025). 10.1161/jaha.124.038945

69 Acoba, M. G. et al. The mitochondrial carrier SFXN1 is critical for complex III integrity and cellular metabolism. Cell reports 34, 108869 (2021). 10.1016/j.celrep.2021.108869

70 Chen, E. Y. et al. Enrichr: interactive and collaborative HTML5 gene list enrichment analysis tool. BMC Bioinformatics 14, 128 (2013). 10.1186/1471-2105-14-128

71 Hao, Y. et al. Dictionary learning for integrative, multimodal and scalable single-cell analysis. Nat Biotechnol 42, 293–304 (2024). 10.1038/s41587-023-01767-y

72 Yu, F., Haynes, S. E. & Nesvizhskii, A. I. IonQuant Enables Accurate and Sensitive Label-Free Quantification With FDR-Controlled Match-Between-Runs. Mol Cell Proteomics 20, 100077 (2021). 10.1016/j.mcpro.2021.100077

