## Supplemental Figures for "Stoichiometric m6A regulation of YY1 orchestrates the metabolic switch during cardiac development"

**Supplemental Figure 1**

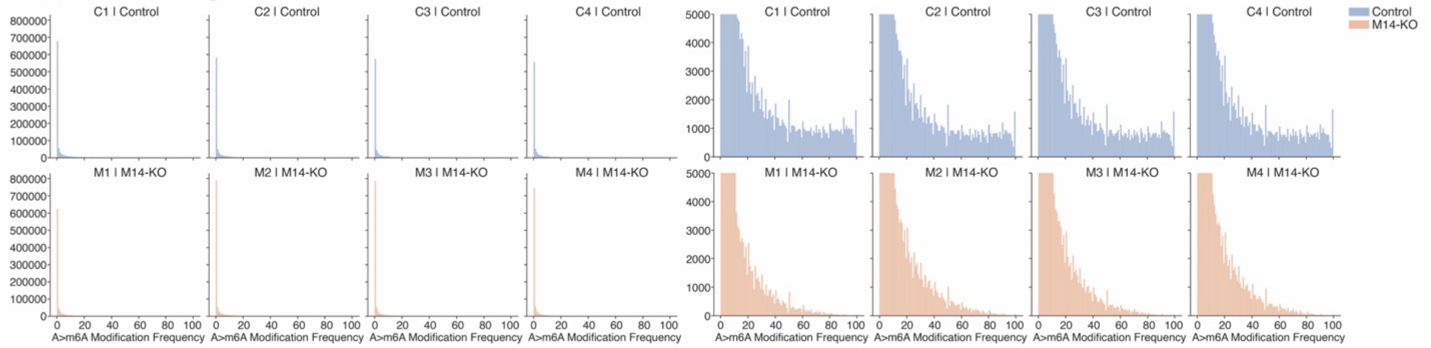


**Supplemental figure S1.** Global decline of m6A modification levels in E12.5 M14-KO hearts.

Histograms showing the distribution of site-level percent m6A modification across all nanopore sequencing replicates (n=4 hearts per condition), and their respective zoom in versions highlighting a global reduction in high-confidence m6A-mdified sites in M14-KO samples compared to controls. Sites with m6A probability ≥0.9 (calculated from modkit pileup) classified as methylated. Only methylation sites with ≥10 reads are represented.

### Supplemental Figure 2

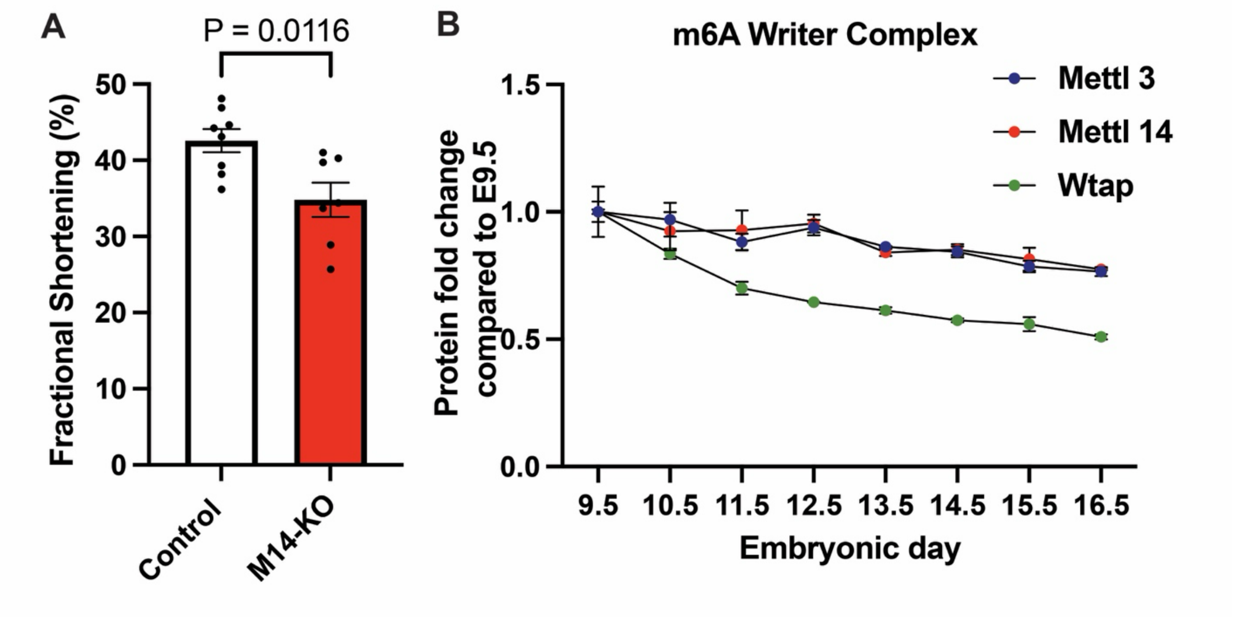


**Supplemental Figure S2.** m6A writer abundance during heart development and cardiac dysfunction in E15.5 M14-KO hearts.

(A) Quantification of left ventricular fractional shortening using in utero echocardiography in E15.5 control (n=8) and M14-KO (n = 7) embryos. M14-KO hearts display reduced left ventricular systolic function compared to controls.

(B) Normalized tandem mass tag-based protein abundances of Mettl3, Mettl14, and Wtap in whole mouse hearts from E9.5 to E16.5 (n=3 embryos per time point; Edwards et al., 2023[1]). Data are shown as mean ± SEM and normalized to E9.5. All three m6A writer proteins are stably expressed around mid-gestation.

All quantitative data are presented as mean ± SEM. The Shapiro-Wilk test was performed to assess normal distribution, and student parametric t-test (two-tailed) was used for all comparisons. Only *P* values <0.1 are reported.

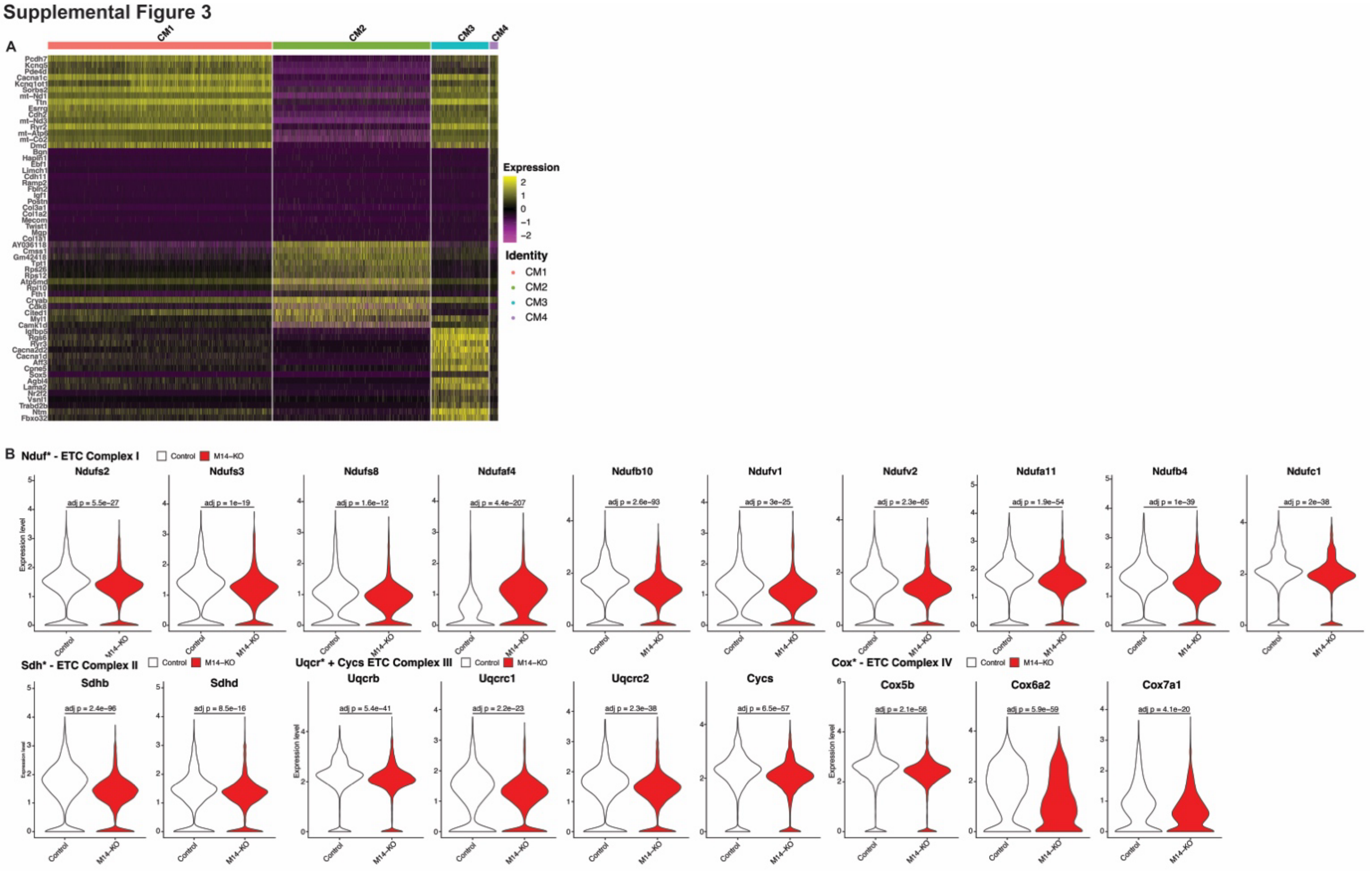


**Supplemental Figure S3.** Cardiomyocyte subtype markers and dysregulated electron transport chain genes in M14-KO hearts.

(A) Heatmap showing the top 15 enriched marker genes for each cardiomyocyte (CM) subpopulation.

(B) Violin plots showing differentially expressed electron transport chain-related genes between Control and M14-KO CMs, grouped by each complex. Gene-level statistics were computed using a two-sided Wilcoxon rank-sum test and adjusted using the Bonferroni method.

### Supplemental Figure 4

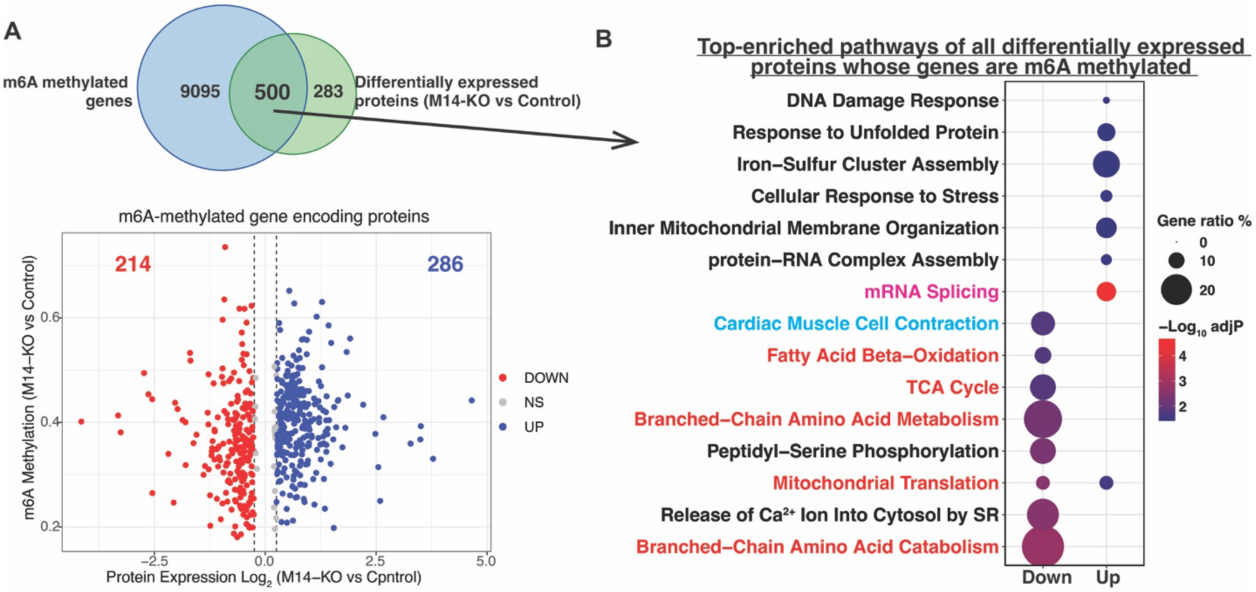


**Supplemental figure S4.** The identified m6A-regulated proteome showed reduced activity in aerobic metabolic pathways.

(A) Integration of m6A modified genes and differentially expressed proteins in M14-KO hearts followed by their respective volcano plot distribution.

(B) GO enrichment analysis of the m6A-modified and differentially expressed proteins revealed a significant association with the downregulation of aerobic metabolic (red) and cardiomyocyte contraction (blue) pathways.

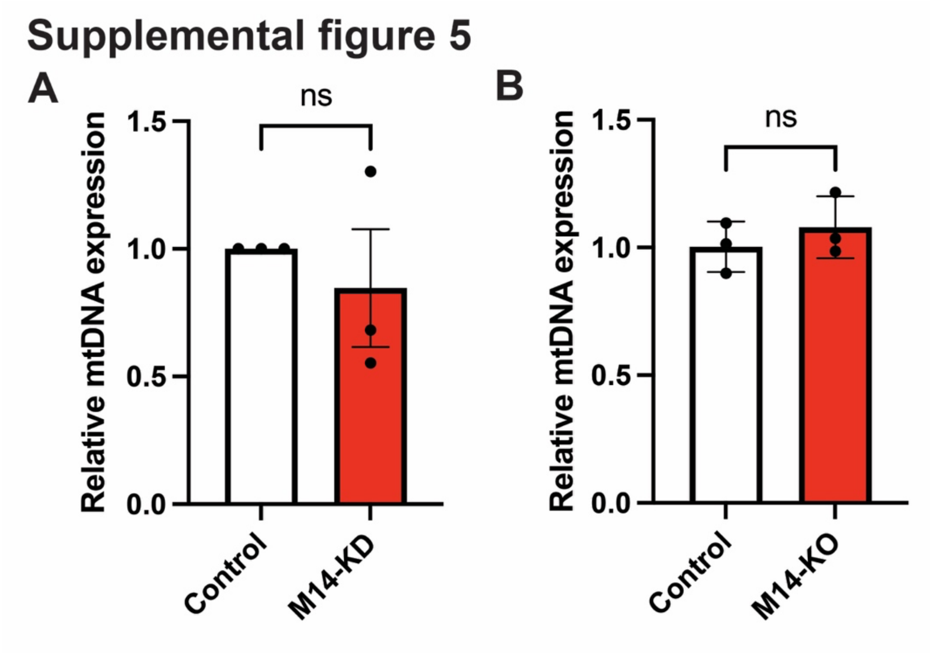


**Supplemental figure S5.** No difference in mitochondrial DNA level (mt-ND1 expression) is observed in both (A) M14-KD hiPSC-CMs (n=3) and (B) M14-KO hearts (n=3). All quantitative data are presented as mean ± SEM. The Shapiro-Wilk test was performed to assess normal distribution, and student parametric t-test (two-tailed) was used for all comparisons. Only *P* values <0.1 are reported.

### Supplemental Figure 6

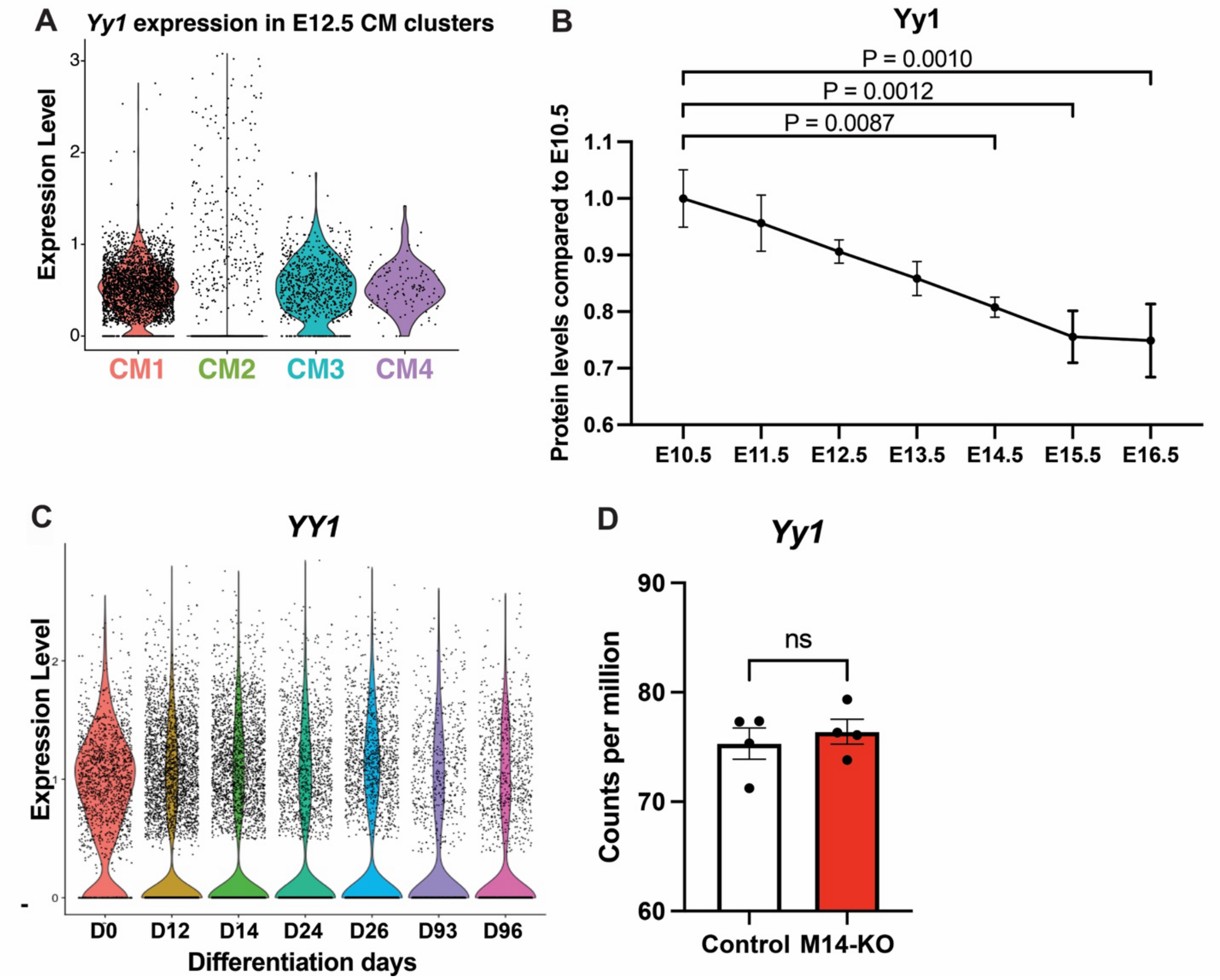


**Supplemental figure S6. A.** *Yy1* gene expression was unchanged in M14-KO E12.5 hearts. **B.** *Yy1* is profoundly downregulated in cluster 2 cardiomyocytes that are characterized by reduced expression of OXPHOS genes. **C.** Proteomics analysis of mouse embryonic hearts shows that Yy1 protein levels continuously decrease throughout mid-gestation[1]. **D.** Single cell analysis of *in vitro* cultured hiPSC-CMs showing stable YY1 expression in hiPSC-CMs that remain persistently immature[2].

All quantitative data are presented as mean ± SEM. The Shapiro-Wilk test was performed to assess normal distribution, and student parametric t-test (two-tailed), Mann-Whitney (two-tailed) non-parametric tests or one-way ANOVA with Bonferroni's multiple comparisons test were used as appropriate for all comparisons. For single cell RNA-Seq analysis statistics were performed as described in Methods. Only *P* values <0.1 are reported.

### References


1 Edwards, W. *et al.* Quantitative proteomic profiling identifies global protein network dynamics in murine embryonic heart development. *Developmental cell* **58**, 1087-1105 e1084 (2023). https://doi.org:10.1016/j.devcel.2023.04.011

2 Kannan, S. *et al.* Trajectory reconstruction identifies dysregulation of perinatal maturation programs in pluripotent stem cell-derived cardiomyocytes. *Cell reports* **42**, 112330 (2023). https://doi.org:10.1016/j.celrep.2023.112330
